# Pyro: A Comprehensive Pipeline for Eukaryotic Genome Assembly

**DOI:** 10.1101/2023.04.18.537425

**Authors:** Dean Southwood, Rahul V. Rane, Siu Fai Lee, John G. Oakeshott, Shoba Ranganathan

## Abstract

The assembly of reference-quality, chromosome-level genomes for both model and novel eukaryotic organisms is an increasingly achievable task for single research teams. However, the broad variety of sequencing technologies, assembly algorithms, and post-assembly processing tools currently available means that there is no clear consensus on a best-practice computational protocol for eukaryotic *de novo* genome assembly. An ever-increasing field of algorithms and packages with unique parameters, setup requirements, and environments makes it difficult for groups to pick up and test new tools, despite potential benefits. Here, we present a comprehensive Snakemake-based pipeline for eukaryotic genome assembly, *Pyro*, to further assist future *de novo* assembly and benchmarking projects. *Pyro* combines 20 assembly and eight polishing packages, comprising 30 different assembly approaches and up to 48 different polishing approaches in combination. These are available across Illumina short-read, Nanopore and PacBio CLR long-read technologies in one container, complete with data preparation, quality metric calculation and result reporting. We demonstrate *Pyro* effectiveness by running *Pyro* on publicly available Illumina, Nanopore and PacBio CLR read sets for *Arabidopsis thaliana*, producing 12 candidate assembly options with minimal initial input or configuration, each with extremely high contiguity and completeness. *Pyro* is highly customizable to expert needs, while also providing an accessible suggested set of tools for more casual users based on simple inputs. *Pyro* is available as a Singularity container suitable for execution on any Unix-compatible OS, and is freely available on GitHub (https://github.com/genomeassembler/pyro). This pipeline provides a one-stop solution for a variety of *de novo* eukaryotic genome assembly needs, and will also assist in the assessment of new tools as a convenient benchmark-generating platform.

## 2 Introduction

The last two decades have seen an explosion of genomics data, with the development of new sequencing technologies and algorithms (Giani et al., 2020), and tools for pre- and post-processing (Dohm et al., 2020). This has created a competitive field of options for those wishing to assemble a high-quality *de novo* genome, highlighted by the variety of methods and approaches used in recent publications (Jayakumar and Sakakibara, 2019; Logsdon et al., 2020).

Next-generation and third-generation sequencing data both play significant roles in current genome assembly practices, particularly when considering the *de novo* assembly of large eukaryotic genomes with extensive repeats. Illumina short-read sequencing provides high per-base-quality reads with low rates of error (Goodwin et al., 2016), but read sizes on the order of 100-300 bases are often insufficient to fully resolve long repeats accurately, even with paired-end sequencing (Tan et al., 2019). Oxford Nanopore long-read sequencing provides ultra-long reads on the order of tens to hundreds of thousands of bases (Jain et al., 2018b), but individual bases have a significantly higher error rate than Illumina reads (Laver et al., 2015). However, Oxford Nanopore reads are relatively cheap - current market prices put them at approximately 250% the price of Illumina, per Gbp. PacBio continuous long read (CLR) sequencing reads provide a compromise, with significantly longer reads than Illumina, but less than Nanopore, and higher per-base quality than Nanopore, but less than Illumina (Amarasinghe et al., 2020). However, these come with a higher price approximately 350% the price of Illumina, per Gbp. The introduction of PacBio circular consensus sequencing (CCS), also known as PacBio HiFi, has provided a long read, high accuracy alternative (Wenger et al., 2019), but for a significant price tag – approximately 1350% the price of Illumina, per Gbp. Given these differences, tools appropriate for each sequencing data type have been tailored to particular features, promoting the development of a wide library of algorithms and packages currently available, each with their own parameters, terminology, dependencies, and runtime environments. Each sequencing technology also requires its own tailored pre- and post-assembly steps such as error correction and polishing, further adding to the number of tools available.

It is difficult to know what assembly package will work best on a given genome ahead of time, and the best assembler for a task will often vary depending on the features of the genome at hand, as well as the sequencing data type selected for the study (Wick and Holt, 2020). However, given the complexity of the field of options and the layers of troubleshooting required when picking up a new tool, there can be a tendency for labs with one reliable assembly package installation to use it for all tasks, regardless of possible applicable options. Unique installation and input requirements for every assembler inevitably dampens uptake of new tools by the wider community, and new users to tools often require time to discover efficient and scalable ways of using them on their own computing environment (Mangul et al., 2019).

Workflow management systems can help linearise inputs and facilitate scale-up of pipelines. Current popular options for bioinformatic purposes vary from programming languages such as CWL (Amstutz et al., 2016), to behind-the-scenes engines such as Cromwell (Voss et al., 2017), to more self-contained systems such as Nextflow (Di Tommaso et al., 2017), Toil (Vivian et al., 2017), Rabix (Kaushik et al., 2017), and Snakemake (Köster and Rahmann, 2012). The field of pipeline frameworks has been extensively reviewed in recent work (Leipzig, 2017). There is active development of pipelines for Nanopore (de Lannoy et al., 2019; Giesselmann et al., 2019; Cozzuto et al., 2020), and PacBio and Illumina (Korhonen et al., 2019) sequencing data, but to our knowledge no pipelines have extensive inclusion across all three technologies for genome assembly, despite their unique benefits (Southwood et al., 2020, unpublished results), and despite groups increasingly sequencing across platforms (Rupp et al., 2018; De Maio et al., 2019).

Here, we provide a comprehensive, scalable, and parallelisable pipeline for *de novo* assembly and polishing of genomes. The pipeline, called *Pyro*, incorporates 20 different assembly packages across three sequencing types, and eight polishing packages across both short and long reads. The pipeline is completely customisable for expert users, with input parameters across tools condensed into common settings. *Pyro* also recommends algorithms for casual users based on input parameters such as estimated genome size, sequencing data types, and computational resources available. We demonstrate the simplicity and effectiveness of *Pyro* by processing publicly available read sets of Illumina, Nanopore, and PacBio data for the model organism the thale cress, *Arabidopsis thaliana* (120 Mbp, approx. 36.7% repeats, five diploid chromosomes), producing 12 final candidate assemblies with high contiguity and high completeness. We expect our pipeline will encourage greater experimentation with, and uptake of, a wider variety of genome assembly tools in the community, as well as greatly facilitate the benchmarking of new assembly and polishing tools against currently available methods on standard datasets.

## 3 Method

### 3.1 Underlying Computational Structures

Given the increasing availability of extensive, high-quality genomic sequencing data, it is more important than ever that computational solutions which handle that data be efficient, parallelisable, and scalable. However, this must not come at the cost of reliability or validity. The *Pyro* pipeline manages this balance by relying primarily on two underlying computational workhorses: the workflow management system Snakemake, and the containerization system Singularity (Kurtzer et al., 2017).

In terms of workflow management systems, Snakemake and Nextflow stand out as flexible options with significant communities, documentation, and integrated support for useful additional features such as high-performance computing (HPC) scheduling systems, and containers. Both options present efficient solutions which scale well given sufficient computational resources. Here, we have chosen to build *Pyro* with Snakemake due to its pythonic nature, ease of use, and ability to be customised with simple additional Python code.

However, given the extensive library of genome assembly options currently available to the community, setting up so many unique systems and environments continues to present a challenge, both in terms of troubleshooting as well as in version consistency. To reduce setup overhead for users, and to increase reliability and reproducibility, we have chosen to containerize all dependencies and components required for *Pyro* using Singularity. Current popular tools for containerization are Singularity and Docker (Merkel, 2014). Docker has an arguably larger library of base packages and Docker containers at present, while Singularity is a newer solution to the market. However, the main differences between the two options are the privileges and permissions required to run each. Docker requires root privileges, presenting potential security issues for flawed or malicious code, and creating potential issues on HPC systems, while Singularity only requires user-level privileges when being run user-side, with root privileges only required during development, a process which can be done pre-distribution (Kurtzer et al., 2017). Due to the extensive reliance at present on HPC systems for genomics researchers, we have elected to containerise the dependencies and components of *Pyro* with Singularity.

Taken together, Snakemake and Singularity allow *Pyro* to be highly efficient, parallelisable, and scalable, as well as flexible, reliable, and reproducible in distribution, even on very different computing environments. *Pyro* is easy to pick up for new users, while maintaining power and flexibility for experienced players in the field, and comes with thorough documentation following best practice guidelines (Karimzadeh and Hoffman, 2018).

### 3.2 Overview of the *Pyro* Pipeline

The *Pyro* pipeline allows users to take raw sequencing data from Illumina short reads or PacBio CLR or Oxford Nanopore long reads, and produce polished candidate assemblies, in a completely automated manner if desired, using either user-selected or pipeline-recommended assemblers and polishers. It also calculates common desired assembly metrics for all assemblies, to allow users to compare the candidates in a meaningful, straightforward manner. In addition, each module can be used independently to process other data supplied by the user, presenting a modular solution that is easily integrable into other preferred method workflows (**Figure 1**). However, if other intermediate programs are not required or desired, it is recommended to run the entire *Pyro* pipeline from start to end as this will make sure common formatting issues for particular packages are resolved, for example whether the input data should be interleaved, have particular header formats, or be compressed or uncompressed. The pipeline consists of four modules:

**FIGURE 1.**
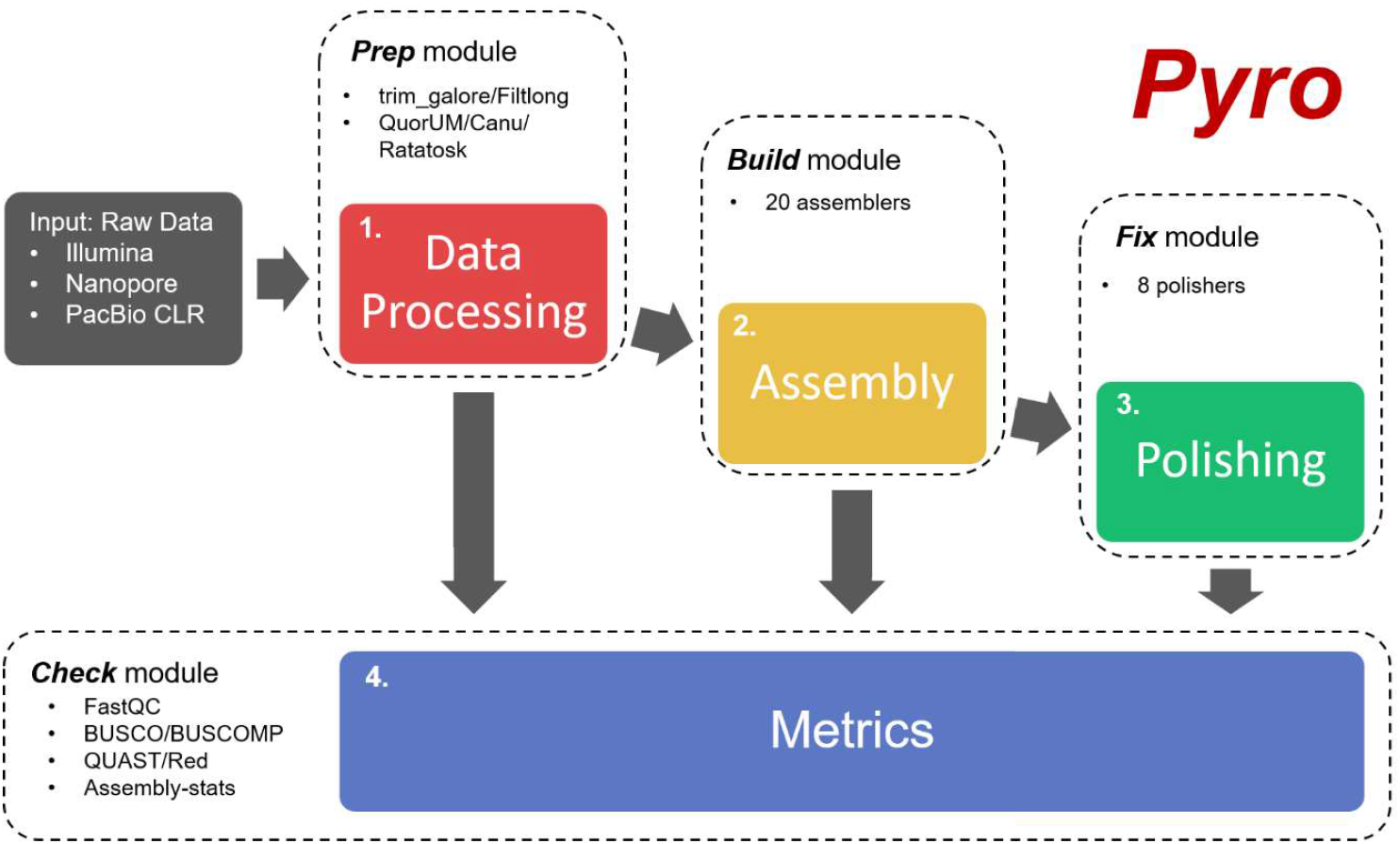
Overview of the *Pyro* pipeline for the construction of high-quality eukaryotic genomes. The pipeline takes raw reads from any combination of Illumina paired-end short reads, Oxford Nanopore long reads, and PacBio CLR technologies. By default, the input reads are filtered and trimmed for quality and adapter sequences using the *Prep* module, and a report of read quality before and after is prepared using the *Check* module. Next, the reads are assembled using the *Build* module, using the three most likely high-performing assemblers by default, based on estimated genome size, read coverage, and computational resources available, with statistics calculated based on the raw assembly using the *Check* module. Finally, the raw assemblies are polished with available reads using the *Fix* module, with final assemblies compared across standard contiguity and gene completeness metrics. All steps are highly customisable to use as few or as many packages as desired.

i. *Prep*: using raw Illumina, PacBio CLR or Oxford Nanopore data as input, this module performs low-level quality control to remove adapters, ambiguous bases, and low-quality reads before assembly. Upon user request, this module will also error-correct the supplied reads if desired.
ii. *Build*: using pre-processed reads from the *Prep* module, or as supplied by user, this module assembles the reads using the selected or recommended assembler(s), outputting draft FASTA assembly files.
iii. *Fix*: using the draft assembly file(s) from the *Build* module, or as supplied by user, this module polishes each draft assembly using the selected or recommended polisher(s) for a selected or recommended number of iterations, depending on input, outputting polished FASTA assembly files.
iv. *Check*: using input from either the *Prep, Build*, or *Fix* modules, or as supplied by user, this module performs metric calculations on the reads and/or assembly files for common desired metrics, including read quality, assembly contiguity, gene completeness, and repeat content. If a reference genome is supplied, this module will also run reference-based comparison calculations to determine genome fraction, as well as rates of common errors such as misassemblies, mismatches, and insertions/deletions (indels), while also providing basic dot plots aligning each candidate assembly to the reference.

### 3.3 Data Preparation Module: *Prep*

The first module of the *Pyro* pipeline, *Prep*, focusses on preparing data for input into genome assembly algorithms. The steps in this preparation depend on the data supplied to the pipeline, as well as what quality control measures are desired by the user. By default, the *Prep* module performs adapter trimming and quality filtering of Illumina reads using the trim_galore package,^1^ and trimming and quality filtering of Oxford Nanopore and PacBio CLR sequences using the Filtlong package.^2^ In addition to this, the *Prep* module will also format the input read files as necessary for use in assemblers by modifying headers, uncompressing files, or interleaving reads as required.

Two additional features of the *Prep* module are not run by default, but are likely to be common desired processes. First, *Prep* is able to perform read correction on short or long reads, using a variety of packages, as can be selected by the user (**Table 1**). While our experience with the assemblers and polishing algorithms available in *Pyro* is that they do not necessitate prior error correction beyond quality filtering to achieve good results, we acknowledge this may not hold true for all genomes or input data. If error-corrected data is to be used downstream for assembly and/or polishing, the --use-corrected flag can be set in subsequent modules. In addition to error correction, the *Prep* module can subsample the supplied input reads to a desired level of coverage using the reformat.sh component of the BBMap/BBTools package (Bushnell, 2014),^3^ based on random seeds, to create smaller coverage for downstream purposes, for example when benchmarking additional tools, or when memory is at a premium.

**TABLE 1.**
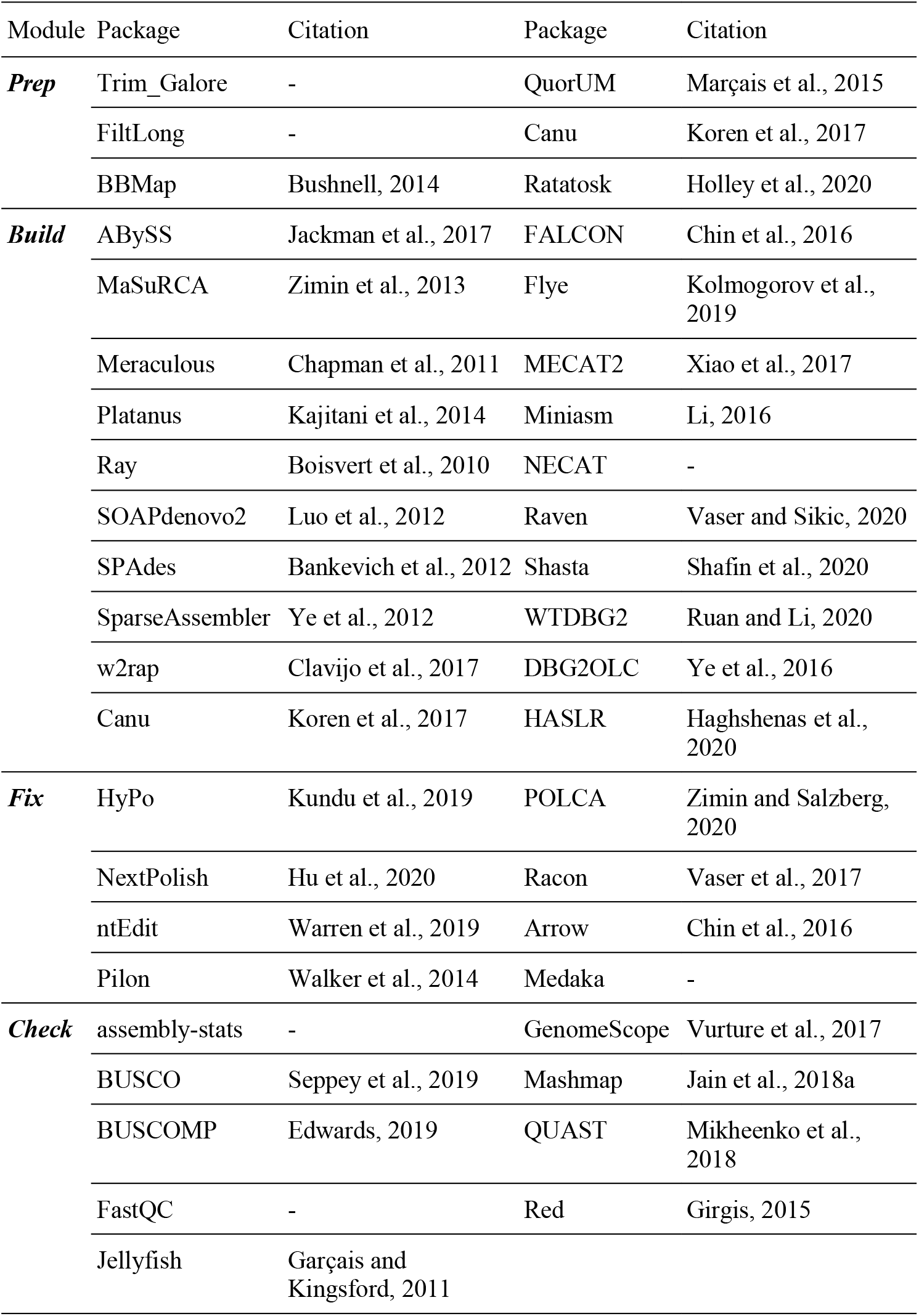
The third-party bioinformatic packages included in the *Prep, Build, Fix*, and *Check* modules of the *Pyro* pipeline. The details and default parameters for each package are available in **Supplementary Tables S1 S4**.

| Module | Package | Citation | Package | Citation |
| --- | --- | --- | --- | --- |
| **Prep** | Trim_Galore | - | QuorUM | Marçais et al., 2015 |
|  | FiltLong | - | Canu | Koren et al., 2017 |
|  | BBMap | Bushnell, 2014 | Ratatosk | Holley et al., 2020 |
| **Build** | ABYSS | Jackman et al., 2017 | FALCON | Chin et al., 2016 |
|  | MaSuRCA | Zimin et al., 2013 | Flye | Kolmogorov et al., 2019 |
|  | Meraculous | Chapman et al., 2011 | MECAT2 | Xiao et al., 2017 |
|  | Platanus | Kajitani et al., 2014 | Miniasm | Li, 2016 |
|  | Ray | Boisvert et al., 2010 | NECAT | - |
|  | SOAPdenovo2 | Luo et al., 2012 | Raven | Vaser and Sikic, 2020 |
|  | SPAdes | Bankevich et al., 2012 | Shasta | Shafin et al., 2020 |
|  | SparseAssembler | Ye et al., 2012 | WTDBG2 | Ruan and Li, 2020 |
|  | w2rap | Clavijo et al., 2017 | DBG2OLC | Ye et al., 2016 |
|  | Canu | Koren et al., 2017 | HASLR | Haghshenas et al., 2020 |
| **Fix** | HyPo | Kundu et al., 2019 | POLCA | Zimin and Salzberg, 2020 |
|  | NextPolish | Hu et al., 2020 | Racon | Vaser et al., 2017 |
|  | ntEdit | Warren et al., 2019 | Arrow | Chin et al., 2016 |
|  | Pilon | Walker et al., 2014 | Medaka | - |
| **Check** | assembly-stats | - | GenomeScope | Vurtture et al., 2017 |
|  | BUSCO | Sepey et al., 2019 | Mashmap | Jain et al., 2018a |
|  | BUSCOMP | Edwards, 2019 | QUAST | Mikheenko et al., 2018 |
|  | FastQC | - | Red | Girgis, 2015 |
|  | Jellyfish | Garçais and Kingsford, 2011 |  |  |

By default, the *Prep* module additionally calls the *Check* module, as discussed below, to provide reports on input read quality, both before and after any quality control, filtering, or correction steps, using the FastQC package.^4^ The FastQC component provides a HTML-formatted report containing information such as per-base quality, GC content, the presence of adapters, and length profiles.

### 3.4 Genome Assembly Module: *Build*

The second module of the *Pyro* pipeline, *Build*, takes input either from the *Prep* module above, or as otherwise supplied by user, to construct a draft genome assembly. The *Build* module can be run with no input aside from reads and information about computational resources, and will produce three assemblies for each data type by default, running assemblers based on resources and internally-estimated read coverage.

However, the *Build* module is completely customisable – the specific assembly packages run, as well as the number of packages run, can be modified using either the --use-assembler command-line flag to use one particular assembler, or using the assemblers setting in the config file to select multiple assemblers. The *Build* module can run 20 different assembly packages natively, depending on input data (**Table 1**), each supplied in the *Pyro* Singularity container with no additional install required. In terms of setting parameters to be supplied to each assembler, by default the *Build* module will use a core set of parameters which should provide a reasonable assembly in most cases (**Supplementary Table S2**). However, the performance of a given assembler on a particular set of input reads is difficult to predict. Therefore, for intermediate users, *Pyro* provides the option of setting common parameters for all assemblers simultaneously in the config file, which is then internally translated into particular flags and parameter settings for each assembler as required. Common adjustments, such as *k*-mer size for de Bruijn graph-based assemblers, can then be performed across multiple assemblers with a single change. For expert users and those familiar with a particular assembler and its parameters, the config file will also accept commands and parameters as a string which will be directly supplied to the assembly package. It is worth noting that such parameters will be supplied to the assembler as-is, and will overwrite any other default options, so should be used with caution.

By default, the *Build* module additionally calls the *Check* module upon completion, as discussed below, to present basic statistics about the assemblies generated, in order for users to decide which assemblies are worth pursuing with further polishing, whether particular parameters have worked as desired, or whether further polishing is required at all. The default components of the *Check* module, which calculate assembly statistics and contiguity metrics using assembly-stats,^5^ GC content, and repeat percentages using Red (Girgis, 2015),^6^ provide a quick overview of useful information; however, if the --check-busco flag is provided, or intermediate-busco parameter is set to True in the config file, the *Check* module additionally runs the BUSCO (Seppey et al., 2019)^7^ and BUSCOMP (Edwards, 2019)^8^ packages to assess gene completeness of assemblies. For this to be run, it requires the additional input of an OrthoDB (Waterhouse et al., 2013) dataset to be specified, either as a local directory location, or by name, in which case it will be downloaded when required.

### 3.5 Polishing Module: *Fix*

The third module of the *Pyro* pipeline, *Fix*, takes as input either a set of assemblies from the *Build* module above, or other draft assemblies as supplied by the user, and produces a set of polished assemblies as output. The polishers selected by default by the *Fix* module depend on the input data types supplied; it is recommended for long read assemblies from Oxford Nanopore or PacBio CLR sequences that a set of short reads from a technology such as Illumina be supplied as well, to enable *Pyro* to perform short-read polishing in addition to polishing with long reads. The *Fix* module can run eight different polishing packages natively, depending on input data (**Table 1**), each supplied in the *Pyro* Singularity container with no additional install required.

For most polishing algorithms, it is often advantageous to polish more than once – this is natively supported in *Pyro*, and can be specified with the --pol-rounds flag, or the polishing-rounds setting in the config file, either overall or for each individual polisher. By default, the *Fix* module runs long-read polishing algorithms for four iterations, and short-read polishing algorithms for three iterations, based on recent benchmarking results (Southwood et al., 2020, unpublished results). For a given assembly, the *Fix* module will by default choose up to two polishing algorithms to run in series – one long-read algorithm for four iterations, followed by one short-read algorithm for three iterations but any combination of algorithms can be supplied through the config file and run for any input assembly, providing intermediate and expert users considerable flexibility to construct their own workflows within *Pyro*. By default, the *Fix* module will call the *Check* module, as detailed below, to provide statistics, quality metrics, and, if a reference is supplied, additional reference-based statistics, for cases where users wish to benchmark methods, or for re-assembly of older reference genomes using long-read technologies, for example.

### 3.6 Assessment and Reporting Module: *Check*

The final module of the *Pyro* pipeline, *Check*, takes either reads, draft assemblies, or polished assemblies as input, and provides common quality metrics and statistics using a variety of popular state-of-the-art packages (**Table 1**). When the entire pipeline is run in an automated default fashion, the *Check* module is called multiple times through the run, providing intermediate outputs that can be checked without having to wait for the entire run to complete. For either input reads, or after the *Prep* module above has run, the *Check* module will provide quality metrics using the FastQC package,^9^ including information about read quality, length, and adapter content. In addition, if requested or if no indication of coverage or genome size is provided, the *Check* module will provide estimates of the genome size and read coverage via *k*-mer counting using Jellyfish (Marçais and Kingsford, 2011)^10^ and GenomeScope (Vurture et al., 2017).^11^ This is often useful information to compare against the size of generated candidate assemblies, but is also necessary in determining which assembly package is likely to produce the best result.

For input draft or polished assemblies, the *Check* module will calculate basic assembly statistics and contiguity metrics using assembly-stats,^12^ repeat content using Red (Girgis, 2015),^13^ and GC content using internal scripts. In addition, if *Pyro* is supplied with a OrthoDB gene set to check against, either as a local directory or by name for automated download, the *Check* module will analyse the presence of benchmarked universal single-copy orthologues (BUSCOs) using the BUSCO package (Seppey et al., 2019),^14^ as well as BUSCOMP (Edwards, 2019),^15^ to assess the gene completeness of the assembly. By using the BUSCOMP package, the input candidate assemblies will be compared to each other, to determine the relative content of BUSCOs present in each assembly; this is particularly useful for checking assemblies pre-polishing, as the BUSCOMP package has less stringent requirements when determining the presence of BUSCO genes, giving a more realistic indication of potential quality post-polishing (Southwood et al., 2020, unpublished results). If a reference assembly is supplied, as may be the case for benchmarking new tools or when re-assembling a previously assembled genome *de novo*, the *Check* module will also provide reference-based statistics using the QUAST-LG package (Mikheenko et al., 2018). The statistics and metrics calculated by the *Check* module are output in human-readable format as a table and a collection of graphs, as well as in CSV format for useful further processing.

### 3.7 Test Case: *Arabidopsis thaliana*

In order to demonstrate the effectiveness of the *Pyro* pipeline, particularly when run with default parameters out-of-the-box, we processed publicly available read sets for the model organism *Arabidopsis thaliana* containing Illumina, PacBio CLR, and Oxford Nanopore reads (Michael et al., 2018) (**Table 2**). These data were used both individually, as well as collectively through hybrid assembly methods. *A. thaliana* has historically been an important model organism for genetic studies, and its genome was the first whole plant genome sequenced (Arabidopsis Genome Initiative, 2000). The current reference genome for *A. thaliana*, TAIR10,^16^ consists of seven structures, namely five diploid chromosome sequences, a mitochondrion sequence, and a chloroplast sequence, for a total genome size of approximately 120 Mbp. The reference genome for *A. thaliana* is actively curated and of high quality, allowing for detailed comparisons to be made between outputs. The *A. thaliana* data were processed through the whole *Pyro* pipeline using default parameters and recommended assemblers and polishers for each technology type, to give an indication of the quality of assembly to be expected out-of-the-box with no curated parameter-setting. The pipeline was run on a single computing node with 20 CPUs and 128 GB of RAM for all steps.

**TABLE 2.** Statistics and availability information about the read sets used to construct the *A. thaliana* candidate assemblies. Coverage was calculated assuming a genome size of 120 Mbp.

| Sequencing Technology | ENA Run Accession No. | Citation | No. Raw Reads | Raw Coverage | Quality-Filtered Coverage |
| --- | --- | --- | --- | --- | --- |
| **Illumina Paired-End** | ERR2173372 | Michael et al., 2018 | 33,683,902 | 70.2 | 66.9 |
| **Oxford Nanopore** | ERR2173373 | Michael et al., 2018 | 300,071 | 28.5 | 25.7 |
| **PacBio CLR** | ERR2173371 | Michael et al., 2018 | 613,080 | 58.3 | 52.5 |

<sup>16</sup> https://www.arabidopsis.org/

## 4 Results

### 4.1 Test Case: *Arabidopsis thaliana*

The *Pyro* pipeline uses any combination of Illumina, Nanopore, and PacBio CLR data, and is particularly tailored for eukaryotic genomes. Due to its base code relying on Snakemake, it is scalable and parallelisable, and its use of Singularity containerisation makes it easy to run on HPC systems. For large genomes or high coverage input data, *Pyro* when run with default parameters makes decisions about which assemblers are likely to scale well with resources, while also producing high-quality outputs. To demonstrate this, the results from the *Pyro* pipeline run on publicly available data for *A. thaliana* are presented below, where *Pyro* has been run on all default parameters, aside from requesting three assemblies from each category of input, namely from Illumina short reads, Oxford Nanopore long reads, PacBio CLR, and hybrid assemblers using a combination of the other three types, in order to display typical results for each input. The final output in this case consists of 12 high quality assemblies which have been polished and assessed against common metrics, primarily assembly contiguity, gene completeness, and reference-based statistics (**Table 3**).

**TABLE 3.**
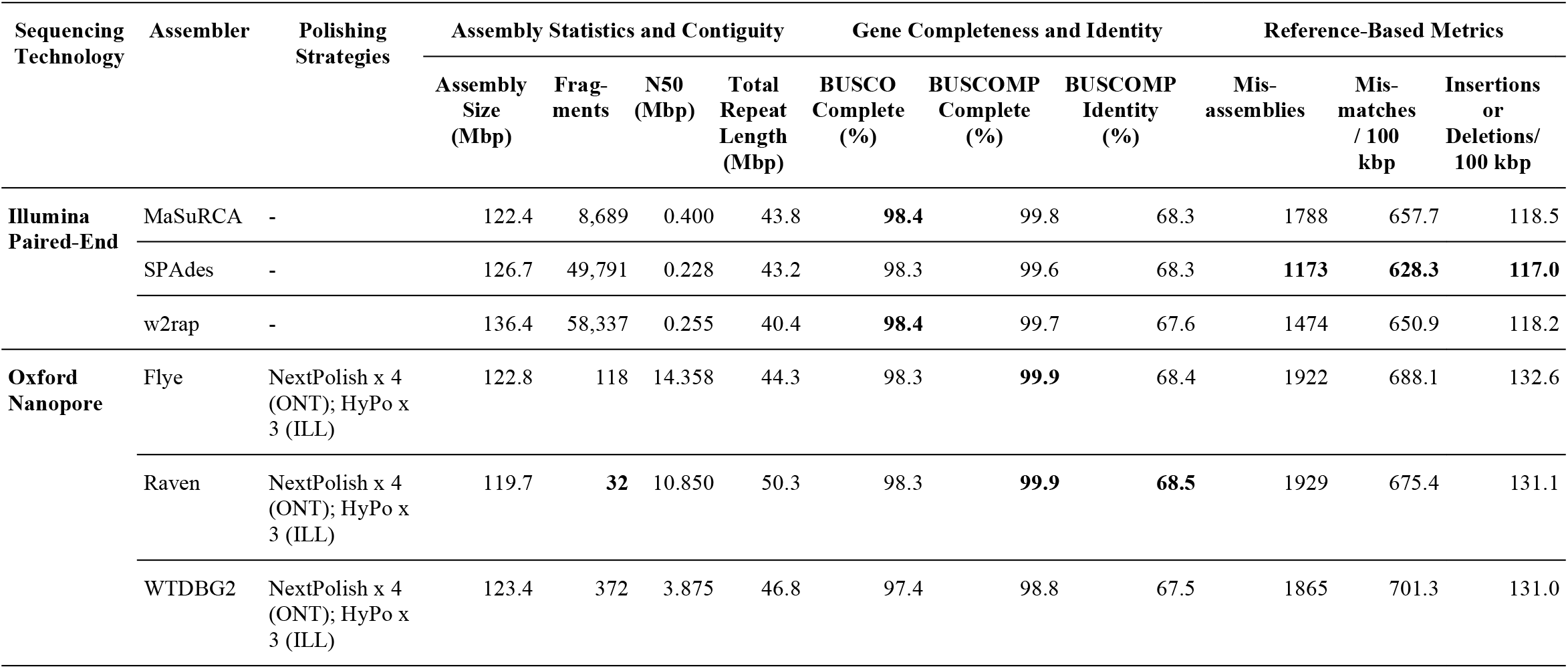
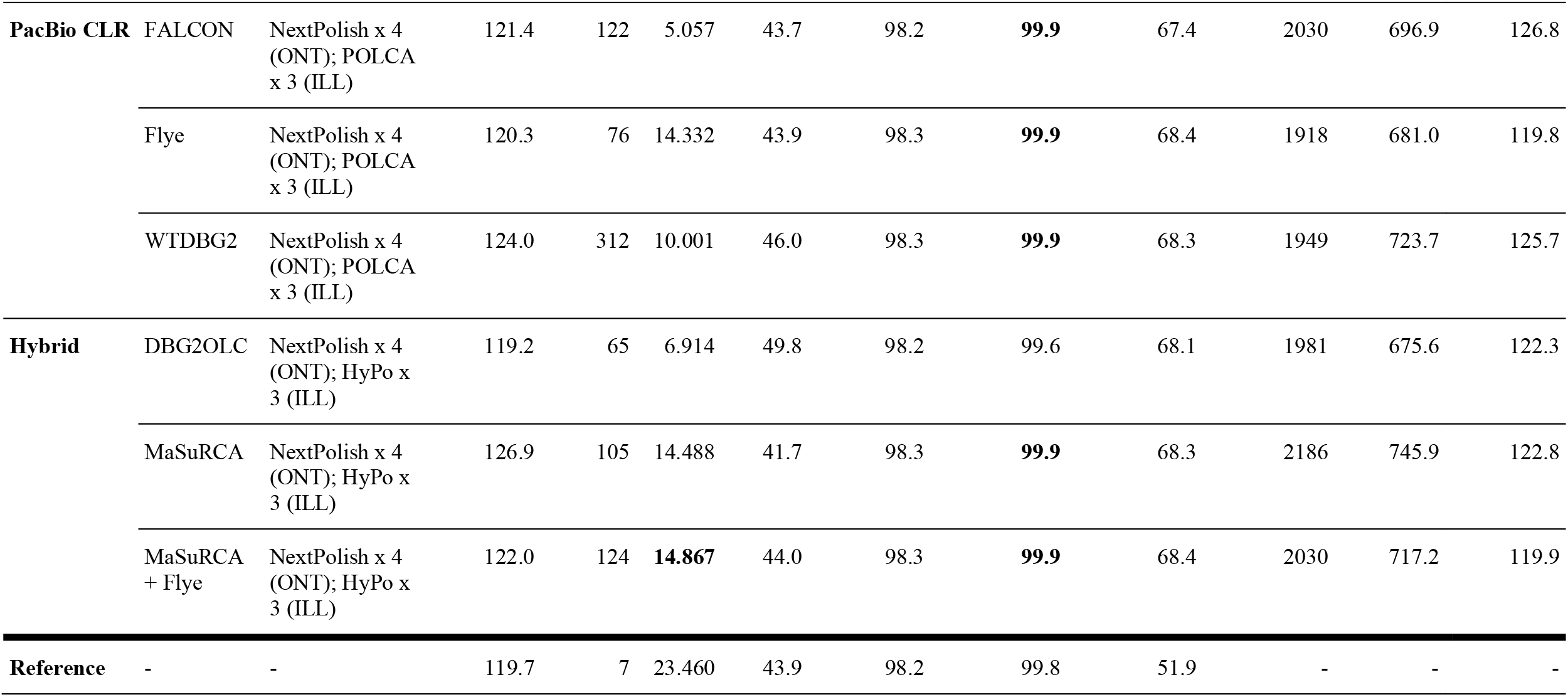
Assembly statistics and quality metric information for the 12 candidate assemblies generated with *Pyro*, highlighting their contiguity and gene completeness, as well as quality metrics calculated against the TAIR10 reference genome. For the Oxford Nanopore, PacBio CLR, and hybrid strategies, the draft assemblies from each assembler were polished with the methods described, and the results for the final polished assemblies are provided; for the Illumina strategies, no polishing was undertaken and the raw assembly statistics are provided. Values in bold are the ‘best’ among all *Pyro* assemblies for metrics where it is useful to rank based on value that is, highest for N50, BUSCO Complete, BUSCOMP Complete, and BUSCOMP Identity, and lowest for Fragments, Misassemblies, Mismatches / 100 kbp and Insertions or Deletions / 100 kbp. Full results including intermediate steps are provided in **Supplementary Table S5**.

| Sequencing Technology | Assembler | Polishing Strategies | Assembly Statistics and Contiguity |  |  |  | Gene Completeness and Identity |  |  | Reference-Based Metrics |  |  |
| --- | --- | --- | --- | --- | --- | --- | --- | --- | --- | --- | --- | --- |
|  |  |  | Assembly Size (Mbp) | Frag-ments | N50 (Mbp) | Total Repeat Length (Mbp) | BUSCO Complete (%) | BUSCOMP Complete (%) | BUSCOMP Identity (%) | Mis-assemblies | Mis-matches / 100 kbp | Insertions or Deletions/ 100 kbp |
| **Illumina Paired-End** | MaSuRCA | - | 122.4 | 8,689 | 0.400 | 43.8 | **98.4** | 99.8 | 68.3 | 1788 | 657.7 | 118.5 |
|  | SPAdes | - | 126.7 | 49,791 | 0.228 | 43.2 | 98.3 | 99.6 | 68.3 | **1173** | **628.3** | **117.0** |
|  | w2rap | - | 136.4 | 58,337 | 0.255 | 40.4 | **98.4** | 99.7 | 67.6 | 1474 | 650.9 | 118.2 |
| **Oxford Nanopore** | Flye | NextPolish x 4 (ONT); HyPo x 3 (ILL) | 122.8 | 118 | 14.358 | 44.3 | 98.3 | **99.9** | 68.4 | 1922 | 688.1 | 132.6 |
|  | Raven | NextPolish x 4 (ONT); HyPo x 3 (ILL) | 119.7 | **32** | 10.850 | 50.3 | 98.3 | **99.9** | **68.5** | 1929 | 675.4 | 131.1 |
|  | WTDBG2 | NextPolish x 4 (ONT); HyPo x 3 (ILL) | 123.4 | 372 | 3.875 | 46.8 | 97.4 | 98.8 | 67.5 | 1865 | 701.3 | 131.0 |
|  |  |  | Assembly Size (Mbp) | Frag-ments | N50 (Mbp) | Total Repeat Length (Mbp) | BUSCO Complete (%) | BUSCOMP Complete (%) | BUSCOMP Identity (%) | Mis-assemblies | Mis-matches / 100 kbp | Insertions or Deletions/ 100 kbp |
| PacBio CLR | FALCON | NextPolish x 4 (ONT); POLCA x 3 (ILL) | 121.4 | 122 | 5.057 | 43.7 | 98.2 | **99.9** | 67.4 | 2030 | 696.9 | 126.8 |
|  | Flye | NextPolish x 4 (ONT); POLCA x 3 (ILL) | 120.3 | 76 | 14.332 | 43.9 | 98.3 | **99.9** | 68.4 | 1918 | 681.0 | 119.8 |
|  | WTDBG2 | NextPolish x 4 (ONT); POLCA x 3 (ILL) | 124.0 | 312 | 10.001 | 46.0 | 98.3 | **99.9** | 68.3 | 1949 | 723.7 | 125.7 |
| Hybrid | DBG2OLC | NextPolish x 4 (ONT); HyPo x 3 (ILL) | 119.2 | 65 | 6.914 | 49.8 | 98.2 | 99.6 | 68.1 | 1981 | 675.6 | 122.3 |
|  | MaSuRCA | NextPolish x 4 (ONT); HyPo x 3 (ILL) | 126.9 | 105 | 14.488 | 41.7 | 98.3 | **99.9** | 68.3 | 2186 | 745.9 | 122.8 |
|  | MaSuRCA + Flye | NextPolish x 4 (ONT); HyPo x 3 (ILL) | 122.0 | 124 | **14.867** | 44.0 | 98.3 | **99.9** | 68.4 | 2030 | 717.2 | 119.9 |
| Reference | - | - | 119.7 | 7 | 23.460 | 43.9 | 98.2 | 99.8 | 51.9 | - | - | - |

### 4.2 2Assembly Statistics and Contiguity

Assembly statistics for each of the polished candidate assemblies are presented in **Table 3**. The candidate assemblies range in size from 119.2 Mbp to 136.4 Mbp, compared to the reference assembly size of 119.7 Mbp, indicating that all assemblers assembled almost all regions of the genome; possible variation in size could be due to the assembly of alternative haplotypes in some regions, or due to variation in length of repeats, particularly around centromeres and telomeres. To investigate this, dot plots were produced by aligning each candidate assembly to the TAIR10 reference assembly using Mashmap (Jain et al., 2018a) (**Supplementary Figures S1 S4**), with the dot plots of the highest quality assemblies from each sequencing type extracted in **Figure 2**. Assemblies differed on the length of heterochromatic centromeric regions, as well as the assembly fragmentation as a whole.

**FIGURE 2.**
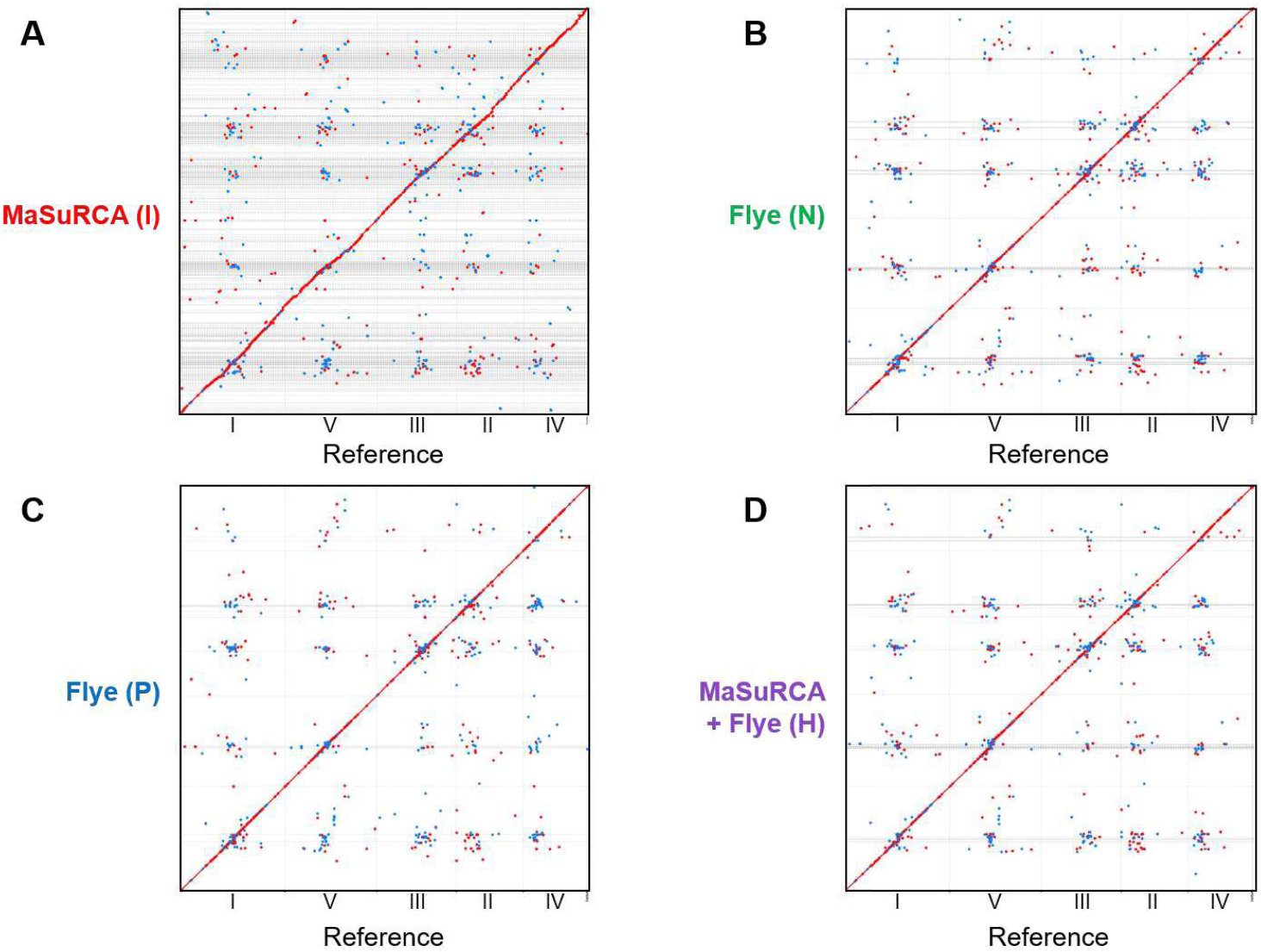
Dot plots generated by aligning the highest quality final candidate assembly of each sequencing type against the TAIR10 reference assembly with Mashmap. Chromosomes in the reference assembly are ordered by total size from largest to smallest. Grey lines indicate contig or scaffold boundaries in each assembly, with higher density lines indicating more fragmented regions. Red dots indicate good alignment between regions of the genome and blue dots indicate weaker alignments, with the substantial red diagonal indicating good reconstruction of the vast majority of the reference. Off-diagonal correlations are largely grouped around heterochromatic regions, likely due to centromeric repeats. (I) = Illumina, (N) = Nanopore, (P) = PacBio, (H) = Hybrid. **(A)** Alignment of the Illumina-only MaSuRCA assembly against the reference, highlighting the high completeness but also high relative fragmentation. **(B)** Alignment of the Oxford Nanopore Flye assembly against the reference, noting the high contiguity and high completeness, with some increased fragmentation around the heterochromatic centromeric regions. **(C)** Alignment of the PacBio CLR Flye assembly against the reference, noting the high contiguity and completeness. **(D)** Alignment of the hybrid MaSuRCA + Flye assembly against the reference, noting the high completeness and contiguity, with some increased fragmentation around the heterochromatic centromeric regions, as in the Oxford Nanopore Flye assembly.

The contiguity of the long-read and hybrid candidate assemblies is consistently high, producing contig N50 lengths as high as 14.8 Mbp. Considering the reference N50 length of 23.5 Mbp, and the lack of any significant scaffolding within the pipeline, this is arguably a very high-quality result, as is also evident from graphically comparing the contiguity with that of the reference (**Figure 3**). The number of contigs for these assemblies was also impressive, with some assembly strategies producing less than 100 contigs, while still covering the vast majority of the reference. For the Illumina-only candidate assemblies, contiguity was considerably lower, as expected given the significantly shorter read length used, and can be seen from the dot plot comparison with the reference assembly (**Supplementary Figure S1**).

**FIGURE 3.**
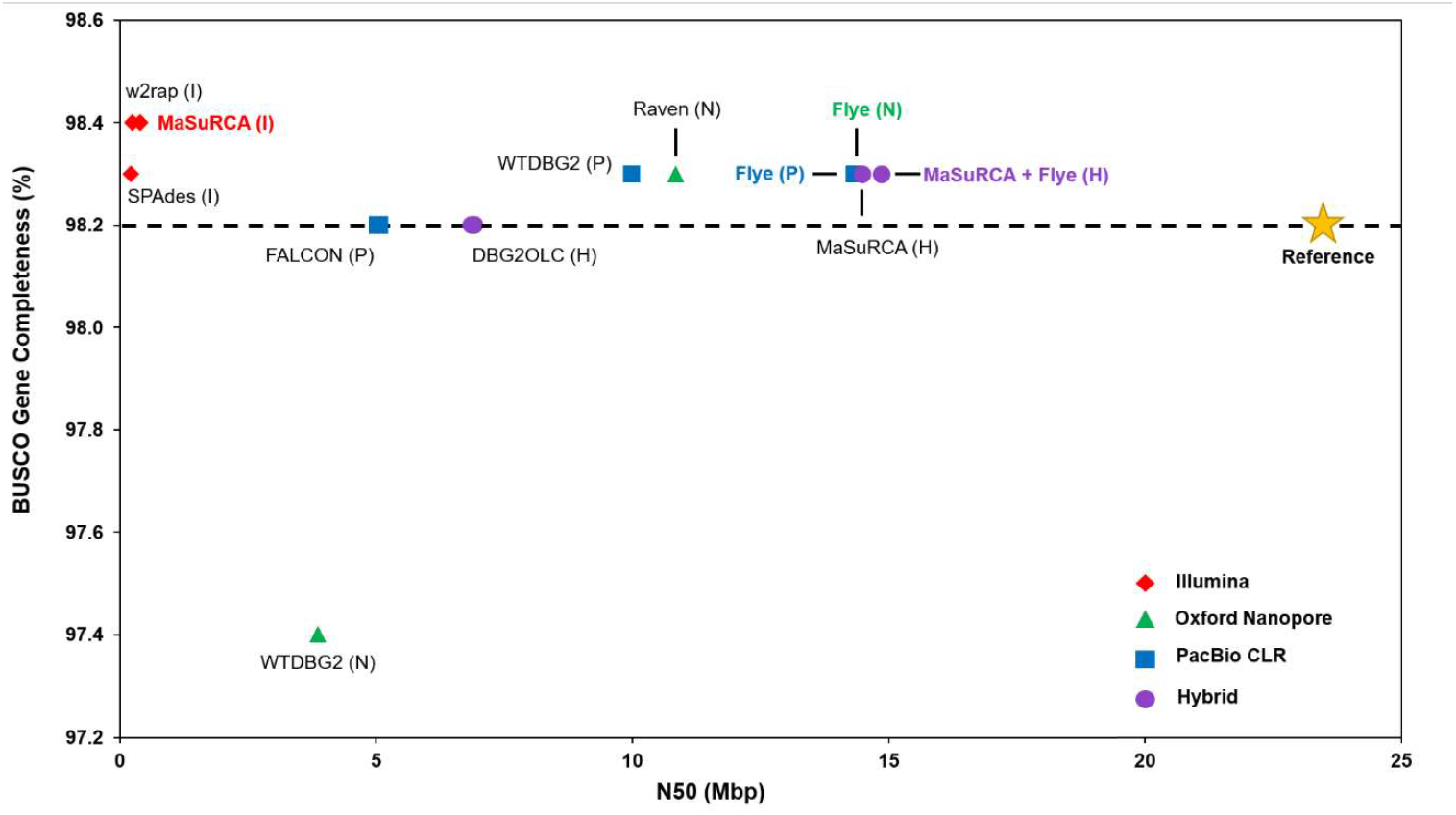
Comparison of contiguity, in terms of N50, and the BUSCO gene completeness of each *A. thaliana* candidate assembly. Each data input type is indicated by shape and colour of markers, as well as in brackets in each label: (I) = Illumina, (N) = Nanopore, (P) = PacBio, (H) = Hybrid. The MaSuRCA (H), MaSuRCA + Flye (H), MaSuRCA (I), SPAdes (I), and w2rap (I) assemblies are in scaffolds, as is the reference; all other assemblies are in contigs. The highest performing assembly from each data input type has a bold, coloured label. The BUSCO gene completeness of the reference assembly, marked as a star, is displayed as a horizontal dotted line.

Taking into consideration the various metrics presented in **Table 3**, we have selected what we consider to be the highest quality assembly from each sequencing type: the MaSuRCA assembly for Illumina paired-end; the Flye assembly for Oxford Nanopore; the Flye assembly for PacBio CLR, and the MaSuRCA + Flye assembly for hybrid sequencing data. These assemblies are shown in bold and coloured font in **Figures 2 6**. Additional results for intermediate steps, such as the raw assemblies and each major stage of polishing, are presented in **Supplementary Table S5**.

### 4.3 Gene Completeness

In order to evaluate the gene completeness of the final assemblies, each candidate assembly was assessed for the presence of BUSCOs using BUSCO v 3 (Seppey et al., 2019) (**Figure 4** and **Table 3**), using the Embryophyta (OrthoDB v 9) gene set consisting of 1440 genes. All twelve candidate assemblies had BUSCO gene completeness within one percent of the reference (98.2%), with over half the candidate assemblies producing a higher BUSCO gene completeness than the reference itself (**Figures 3 4**).

**FIGURE 4.**
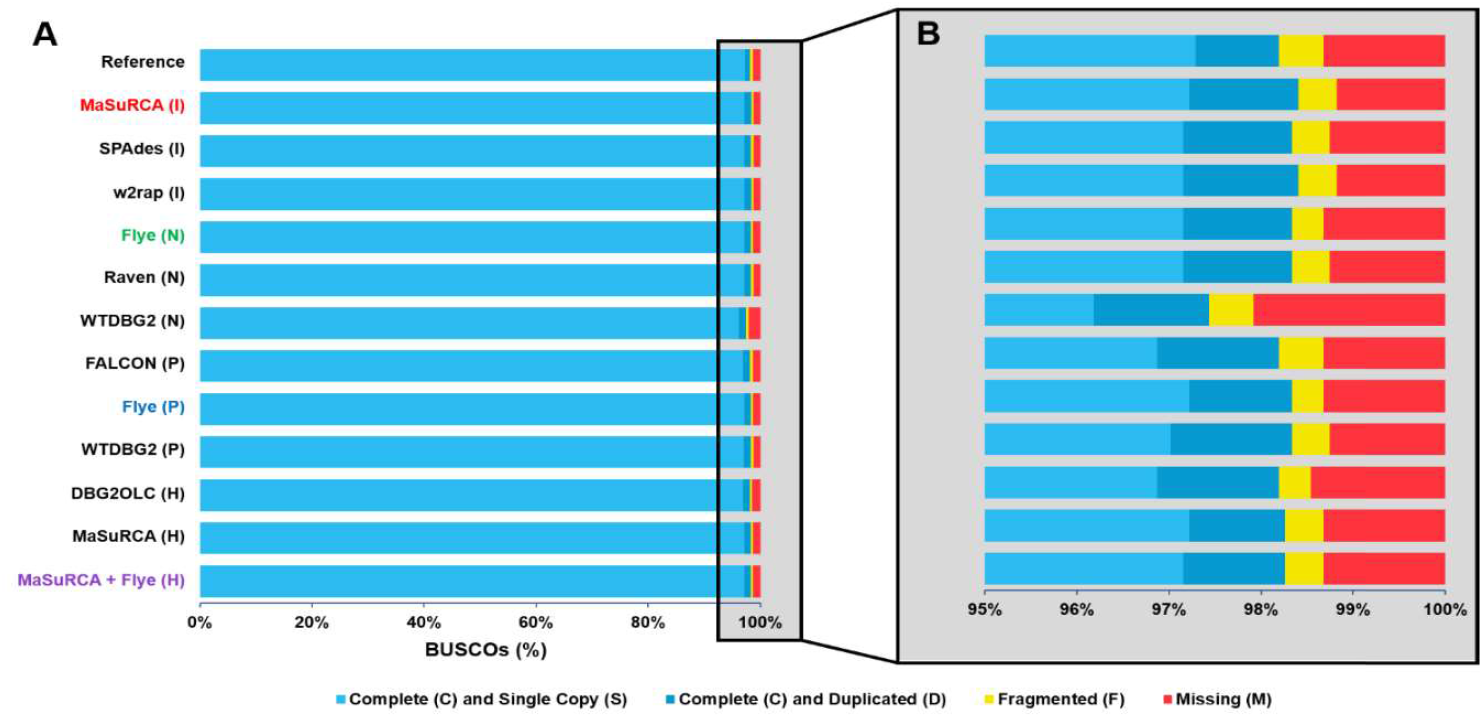
Comparison of gene completeness of all final *A. thaliana* candidate assemblies using BUSCO, analysed against the Embryophyta (OrthoDB v 9) dataset of 1440 genes. (I) = Illumina, (N) = Nanopore, (P) = PacBio, (H) = Hybrid. **(A)** Presence of BUSCO genes in the final candidate assemblies, broken down into complete and single copy genes, complete and duplicated genes, fragmented genes, and missing genes, illustrating the consistently high completeness of all output assemblies. **(B)** Zoom of BUSCO gene plots highlighting the differences between output assemblies.

However, due in part to the stringent thresholds within the BUSCO package, estimation of gene completeness using BUSCO alone often does not show the whole picture (Edwards, 2019; Southwood et al., 2020, unpublished results). For a more comprehensive comparison between the candidate assemblies and the reference, the BUSCO results for all assemblies were re-processed using BUSCOMP (Edwards, 2019), presenting more detailed breakdowns of the presence or absence of BUSCOs. The salient BUSCOMP statistics – namely, the BUSCOMP gene completeness percent, and the BUSCOMP gene identical percent – for each final candidate assembly are presented in **Table 3** and are displayed in standard BUSCO plotted form in **Figure 5**. Using a more relaxed threshold, the completeness is higher when measured by BUSCOMP than by BUSCO, and is more consistent between assemblies, with all but the WTDBG2 Oxford Nanopore assembly having a BUSCOMP gene completeness within 0.2% of the reference. As a useful complement to the more relaxed completeness measure, BUSCOMP also calculates the percentage of sequences in the candidate assemblies which are identical to BUSCO sequences analyzed. The candidate assemblies presented here contain more identical BUSCO genes than the reference, consistently reproducing approximately 68% of genes assessed in identical sequences, compared to 51.9% in the reference. This again speaks in part to the quality of the candidate assemblies across the board, as well as the quality of the data used. Additional results, including for each raw assembler and for the final steps of each method of polishing for each assembler, are displayed in **Supplementary Table S5**.

**FIGURE 5.**
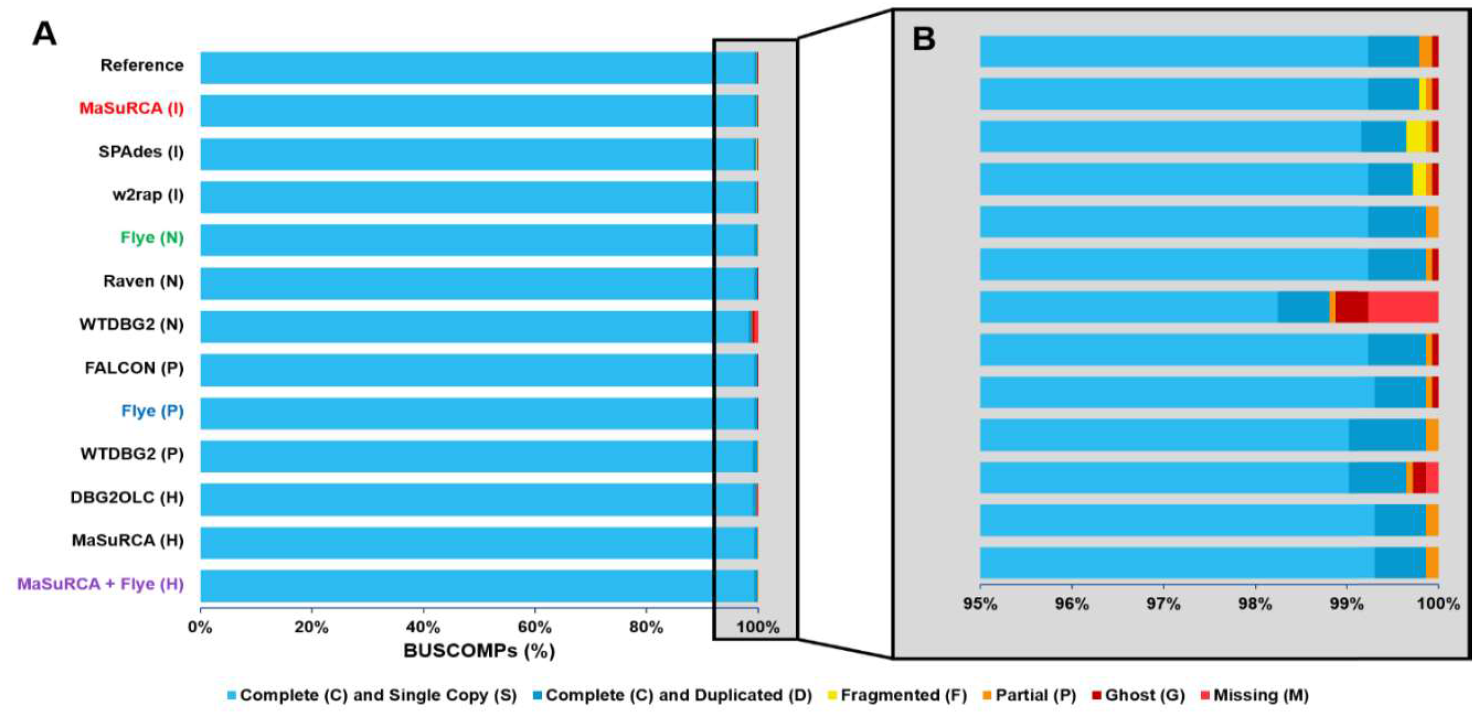
Re-analysis of BUSCO gene completeness of all final *A. thaliana* candidate assemblies using BUSCOMP, with statistics calculated on the 1427 genes present across all candidate assemblies and the reference, out of a possible 1440 BUSCOs analysed. (I) = Illumina, (N) = Nanopore, (P) = PacBio, (H) = Hybrid. **(A)** Presence of BUSCOMP genes in the final candidate assemblies, broken down into complete and single copy genes, complete and duplicated genes, fragmented genes, partial genes, ghost genes, and missing genes, illustrating again the consistently high completeness of all output assemblies. **(B)** Zoom of BUSCOMP gene plots highlighting the differences between output assemblies, noting the higher fragmentation of the Illumina-only assemblies.

### 4.4 Reference-Based Comparisons

In order to characterise the quality of the potential outputs of the pipeline, we have relied on a well characterised model organism with a high-quality reference genome. Given this reference, the *Pyro* pipeline makes use of QUAST v 5.2.0 (Mikheenko et al., 2018) to calculate reference-based statistics such as genome fraction, misassemblies, mismatches, and insertion/deletion errors. The reference-based statistics for each candidate assembly are presented in **Table 3**. In general, all candidate assemblies performed relatively well across all metrics. However, the Illumina-only assemblies provide the lowest number of misassemblies, mismatches, and insertion/deletions, due in part to the low error rate of the Illumina input data. However, given the low contiguity of the Illumina-only assemblies, it is perhaps unsurprising that they achieve the least misassemblies. The Oxford Nanopore-only assemblies tended to produce slightly higher rates of insertion/deletion errors in the final assemblies. However, this is a dramatic reduction from the raw assemblies, and even the assemblies polished additionally with Oxford Nanopore reads alone the Illumina polishing steps in the *Pyro* pipeline reduce the insertion/deletion errors in these assemblies by almost ten-fold (**Supplementary Table S5, Supplementary Figure S5**). This is balanced somewhat by a small increase in mismatches.

A visual indication of the quality of the assemblies generated is illustrated by aligning the final assemblies to the reference, as displayed by dot plots (**Supplementary Figure S1**), with the dot plots for the best assembly from each sequencing type extracted in **Figure 2**. There is a clear increase in contiguity for the long-read and hybrid assembly strategies, but all assemblies construct the vast majority of the reference. In particular, the long-read assembly strategies largely reconstruct the chromosomal arms of the reference genome in single contigs, with more fragmentation evident in the heterochromatic regions of the genome. Of note is the differences in length within the centromeric region of chromosome 5 – this is perhaps unsurprising, given previously reported evidence that it contains large repetitive regions which have historically been difficult to sequence with short-read sequencing alone in prior assembly efforts (Kazusa DNA Research Institute et al., 2000). This is also the case with chromosome 2, although to a lesser extent, but similarly fits with previous sequencing efforts (Lin et al., 1999) Additional accuracy and reference-based statistics for all intermediates are displayed in **Supplementary Table S5**. Additional dot plots for all final assemblies are displayed in **Supplementary Figures S1 S4**.

### 4.5 Computational Performance

The *Pyro* pipeline is capable of efficiently utilising local computational resources, as well as running on HPC systems with minimal configuration overhead. If the --benchmark flag is set, *Pyro* will report run time statistics, such as total run time and maximum memory usage for each assembler, polisher, and metric checking step, providing a convenient method of benchmarking state-of-the-art tools. An example of such a benchmarking output is given in **Figure 6**, displaying the total run time for each final assembly generated, ranging from as little as 35 minutes, to as much as almost 17 hours. A detailed breakdown of wall clock time and maximum memory usage is provided in **Supplementary Table S6**.

**FIGURE 6.**
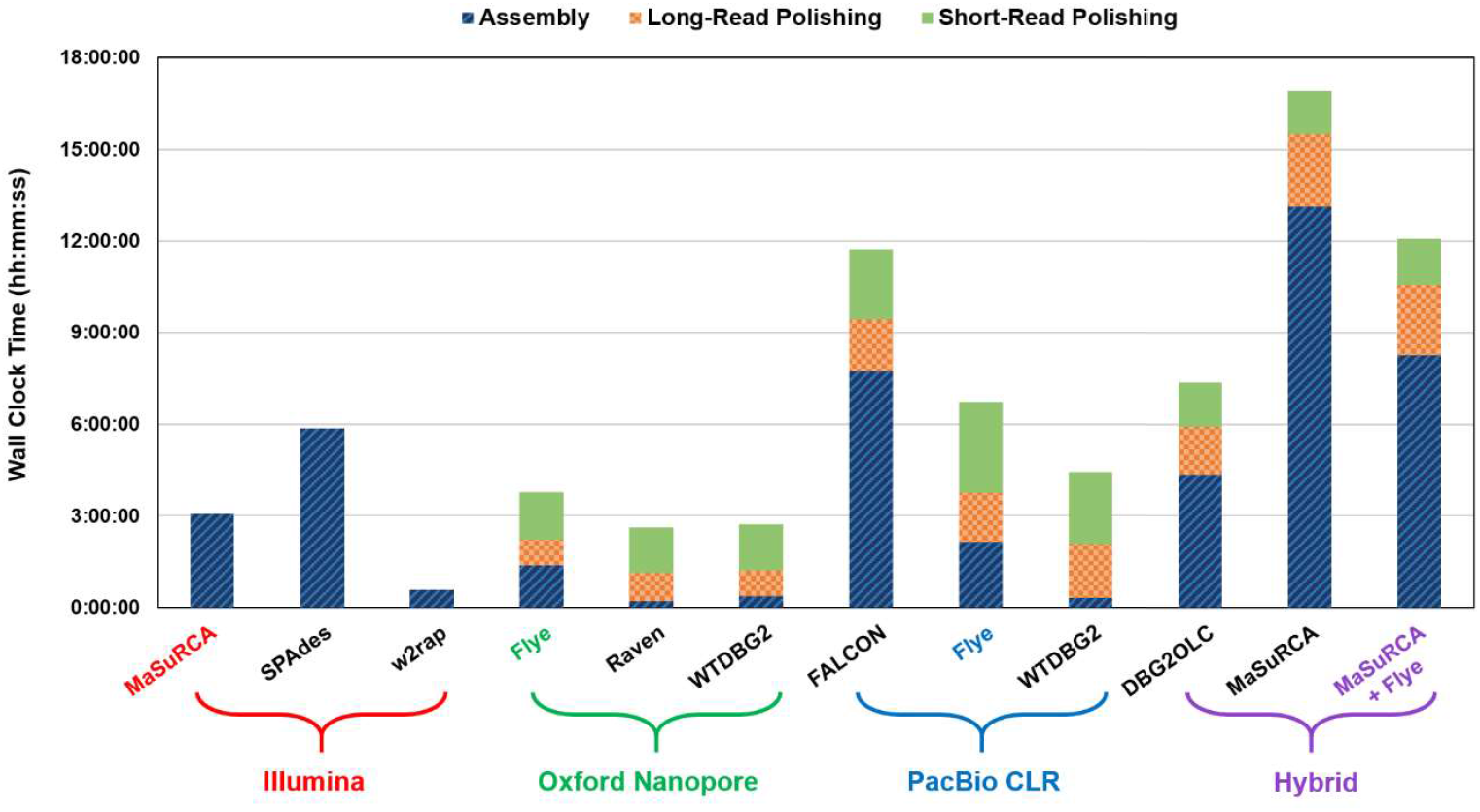
Computational requirements involved for each final polished assembly as measured by wall clock time. For each initial assembler, the time taken by the assembly package is indicated in blue stripes. For the long-read and hybrid assembly strategies, the time taken by the long-read polishing package (NextPolish for all) is indicated in orange checks. Finally, the time taken by the short-read polishing packages (HyPo for Oxford Nanopore and Hybrid, and POLCA for PacBio CLR) is indicated in green dots. The Illumina-only assemblies did not have any polishing done and therefore only the raw assembly time is indicated. The highest quality assemblies for each sequencing type are indicated with colored labels below the plot.

## 5 Discussion

The *Pyro* pipeline presents a scalable, self-contained toolkit for genome assembly. Given input data from most main sequencing platforms, *Pyro* is capable of recommending a suite of assembly and polishing options to suit user-specified needs, and run them, managing all dependencies and parallelisation internally. It is highly customisable to individual needs, and can be tuned via config file to provide consistent user-defined workflow parameters for easy reproducibility. By default, it will select assembly packages likely to perform well using reasonable computational resources, prioritising scalable solutions, particularly for large genomes and high-coverage datasets.

A key strength of the *Pyro* pipeline is in its generation of multiple assembly candidates for a given set of input data with minimal initial configuration. On the *A. thaliana* dataset, *Pyro* produces a dozen high quality candidate assemblies for a variety of downstream purposes, depending on user priorities, whether it be contiguity, gene completeness, or per-base accuracy, along with metrics to help inform decisions about which assembly to pick. These assemblies all covered the majority of the *A. thaliana* reference sequence, including heterochromatic regions (**Supplementary Figure S1**), with as few as hundreds to tens of contigs in the case of long-read assemblers. For most assembly purposes, it will not be necessary to trial 12 assemblers; however, these results indicate that even three assembly options are likely to produce reasonably strong assembly candidates, regardless of input data type.

The *Pyro* pipeline is a large toolkit of assembly and polishing tools, incorporating a wide range of packages across assembly, polishing and evaluation. However, it is still necessarily limited in its scope, and is best supplemented with other post-polishing tools and methods. One key limitation of the pipeline in its current form is the lack of tools specifically tailored to newly developed sequencing data types, such as PacBio circular consensus sequencing (CCS), also referred to as PacBio HiFi, which presents a promising alternative to current long-read technologies (Wenger et al., 2019). However, at present this is still an option with a relatively high price tag. As the results from the test data set indicate, high contiguity, high quality assemblies can be generated from existing technologies for a lower price, presenting more accessible options a key focus of the pipeline. Further post-processing is also possible on the candidate assemblies produced by the pipeline. These can be processed with haplotype reduction tools such as Purge Dups (Guan et al., 2020) or Purge Haplotigs (Roach et al., 2018) to produce single haplotype-resolution assemblies and reduce redundancy.

Polished assemblies can be scaffolded with other sequencing technologies not considered here, such as Hi-C and BioNano sequencing to produce true chromosome-level assemblies, using packages such as ALLHiC (Zhang et al., 2019) and 3D-DNA (Dudchenko et al., 2017). The candidate assemblies produced by *Pyro* can also be used as inputs for assembly merging packages such as Quickmerge (Chakraborty et al., 2016). While these are not natively supported in the current implementation of *Pyro*, the candidate assemblies generated from the pipeline provide robust, high quality input options for these additional methods, reducing the complexity of the genome assembly process, particularly for new researchers to the field.

The portability and ease of use of the *Pyro* pipeline on HPC systems is a key strength. The containerisation of the pipeline presents a highly reproducible option for genome assembly, one which is flexible enough to run on HPC systems without the need for root user privileges. However, this is not without potential drawbacks. In a field with a high state of flux, where new packages are developed and current packages are updated, maintaining a balance between reproducibility and updating is a challenge for all pipelines relying on third-party tools. The use of Singularity presents a potential solution however, allowing for a ‘plug-and-Play‘ container system when significant updates to key packages arise. The Singularity container can be updated and re-released to include the new packages and possible new dependencies, requiring minimal updating or setup by the user, only the exchange of one container for the next, while still maintaining older containers for reproducibility purposes. Striking the balance between these competing interests is essential for any scientific software package development project, with *Pyro* being no exception.

## 6 Availability and Future Directions

The *Pyro* pipeline is coded primarily in Python (3.7), with some scripting in Bash within the Singularity container, and is released under a GNU General Public License v3.0. The pipeline code, installation instructions, and substantial further documentation are publicly available at https://github.com/genomeassembler/pyro, and the pipeline is able to be run on any Linux or Unix operating system capable of running Singularity and Python 3. The pipeline will be kept updated with new assemblers and polishers as they become publicly available, and major updates of currently included assemblers and polishers will be incorporated in further releases.

## 7 Author Contributions

DS wrote the code for the pipeline and ran the test case data. DS, RVR and SFL contributed to code development, optimization, troubleshooting and pipeline features. JGO and SR supervised the project. DS led the writing of the manuscript and made figures and tables, with significant contributions from all authors to the final manuscript.

## 8 Funding

This research has been supported by a Macquarie University Research Training Pathway Scholarship and a Macquarie University Postgraduate Research Funding grant to DS.

## 9 Conflict of Interest

The authors declare that the research was conducted in the absence of any commercial or financial relationships that could be construed as a potential conflict of interest.

## 10. Acknowledgments

We would like to thank the CSIRO Scientific Computing team for assistance in general debugging of features during development.

## Footnotes

1 https://github.com/FelixKrueger/TrimGalore

2 https://github.com/rrwick/Filtlong

3 https://sourceforge.net/projects/bbmap/

4 https://www.bioinformatics.babraham.ac.uk/projects/fastqc/

5 https://github.com/sanger-pathogens/assembly-stats

6 https://github.com/BioinformaticsToolsmith/Red

7 https://busco.ezlab.org/

8 https://github.com/slimsuite/buscomp

9 https://www.bioinformatics.babraham.ac.uk/projects/fastqc/

10 https://github.com/gmarcais/Jellyfish

11 https://github.com/schatzlab/genomescope

12 https://github.com/sanger-pathogens/assembly-stats

13 https://github.com/BioinformaticsToolsmith/Red

14 https://busco.ezlab.org/

15 https://github.com/slimsuite/buscomp

16 https://www.arabidopsis.org/

## Notes

https://github.com/genomeassembler/pyro

### Competing Interest Statement

The authors have declared no competing interest.

