## Supplementary Files for "Pyro: A Comprehensive Pipeline for Eukaryotic Genome Assembly"

Vurture, G. W., Sedlazeck, F. J., Nattestad, M., Underwood, C. J., Fang, H., Gurtowski, J., et al. (2017). GenomeScope: Fast reference-free genome profiling from short reads. *Bionformatics* 33 (14), 2202–2204. doi: 10.1093/bioinformatics/btx153

Walker, B. J., Abeel, T., Shea, T., Priest, M., Abouelliel, A., Sakthikumar, S., et al. (2014). Pilon: An integrated tool for comprehensive microbial variant detection and genome assembly improvement. *PLoS One* 9 (11), e112963. doi: 10.1371/journal.pone.0112963

Warren, R. L., Coombe, L., Mohamadi, H., Zhang, J., Jaquish, B., Isabel, N., et al. (2019). ntEdit: Scalable genome sequencing polishing. *Bioinformatics* 35 (21), 4430–4432. doi: 10.1093/bioinformatics/btz400

Waterhouse, R. M., Tegenfeldt, F., Li, J., Zdobnov, E. M., and Kriventseva, E. V. (2013). OrthoDB: A hierarchical catalog of animal, fungal and bacterial orthologs. *Nuc. Acid Res.* 41 (D1), D358–D365. doi: 10.1093/nar/gks1116

Wenger, A. M., Peluso, P., Rowell, W. J., Chang, P. C., Hall, R. J., Concepcion, G. T., et al. (2019). Accurate circular consensus long-read sequencing improves variant detection and assembly of a human genome. *Nat. Biotech.* 37, 1155–1162. doi: 10.1038/s41587-019-0217-9

Xiao, C. L., Chen, Y., Xie, S. Q., Chen, K. N., Wang, Y., Han, Y., et al. (2017). MECAT: Fast mapping, error correction, and *de novo* assembly for single-molecule sequencing reads. *Nat. Methods* 14, 1072–1074. doi: 10.1038/nmeth.4432

Ye, C., Ma, Z. S., Cannon, C. H., Pop, M., and Yu, D. W. (2012). Exploiting sparseness in *de novo* genome assembly. *BMC Bioinformatics* 13, S1. doi: 10.1186/1471-2105-13-S6-S1

Ye, C., Hill, C. M., Wu, S., Ruan, Y., and Ma, Z. S. (2016). DBG2OLC: Efficient assembly of large genomes using long erroneous reads of the third generation sequencing technologies. *Sci. Rep*. 6, 31900. doi: 10.1038/srep31900

Zhang, X., Zhang, S., Zhao, Q., Ming, R., and Tang, H. (2019). Assembly of allele-aware, chromosomal-scale autopolyploid genomes based on Hi-C data. *Nat. Plants* 5, 833–845. doi: 10.1038/s41477-019-0487-8

Zimin, A. V., Marçais, G., Puiu, D., Roberts, M., Salzberg, S. L., and Yorke, J. A. The MaSuRCA genome assembler. *Bioinformatics* 29 (21), 2669–2677. doi: 10.1093/bioinformatics/btt476

Zimin, A. V., and Salzberg, S. L. (2020). The genome polishing tool POLCA makes fast and accurate corrections in genome assemblies. *PLoS Comput. Biol.* 16 (6), e1007981. doi: 10.1371/journal.pcbi.1007981

#### Data Availability Statement

Sources for sequence data used in this study have been listed in **Table 2**. All results generated or analysed in this study are included in this published article.

### SUPPLEMENTARY MATERIAL

#### 1 Supplementary Figures and Tables

##### 1.1 Supplementary Figures

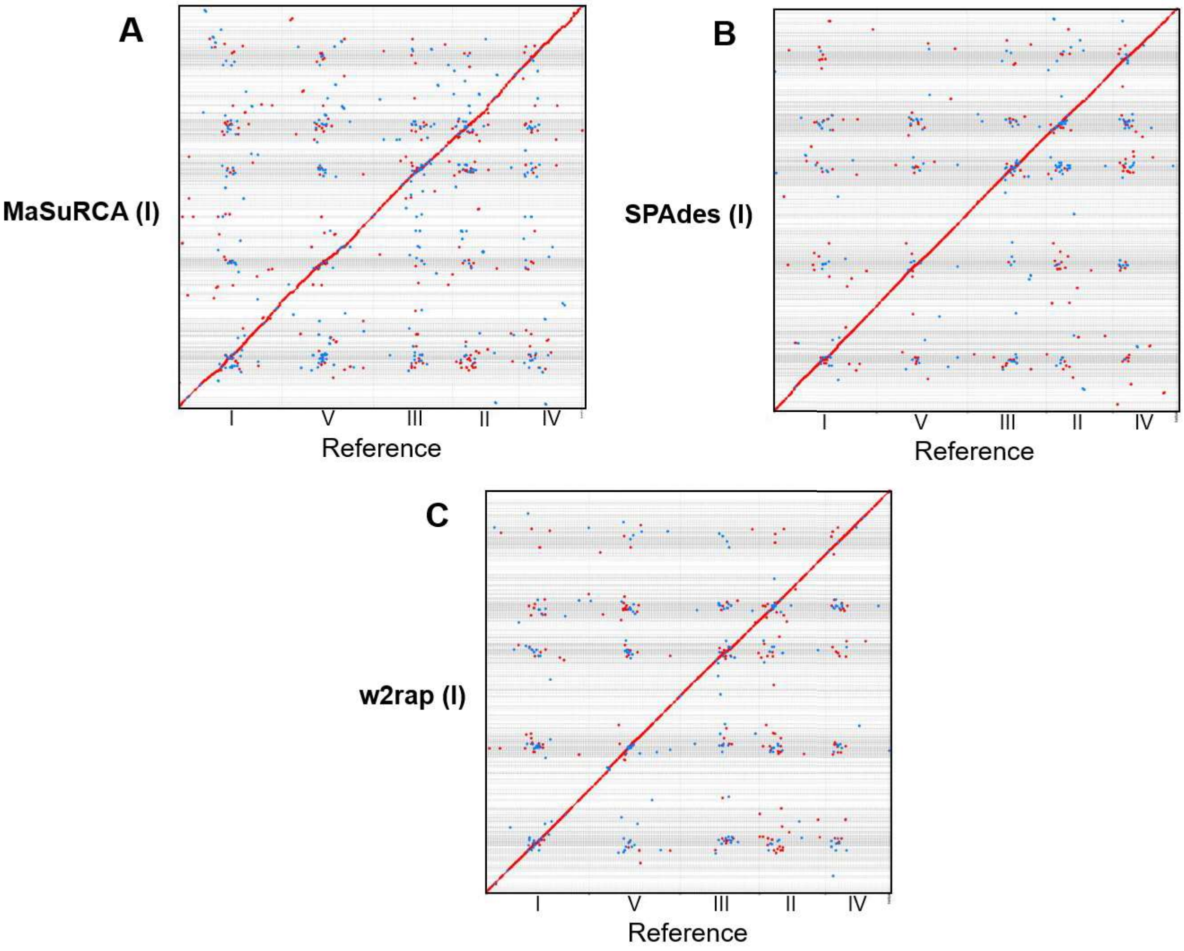


**Supplementary Figure S1 |** Dot plots for all final Illumina-only candidate assemblies generated for *A. thaliana* with the *Pyro* pipeline, as generated by Mashmap in comparison to the TAIR10 reference assembly. (I) = Illumina, (N) = Nanopore, (P) = PacBio, (H) = Hybrid. All assemblies show high completeness when compared with the reference, with expected levels of fragmentation given the short read length.

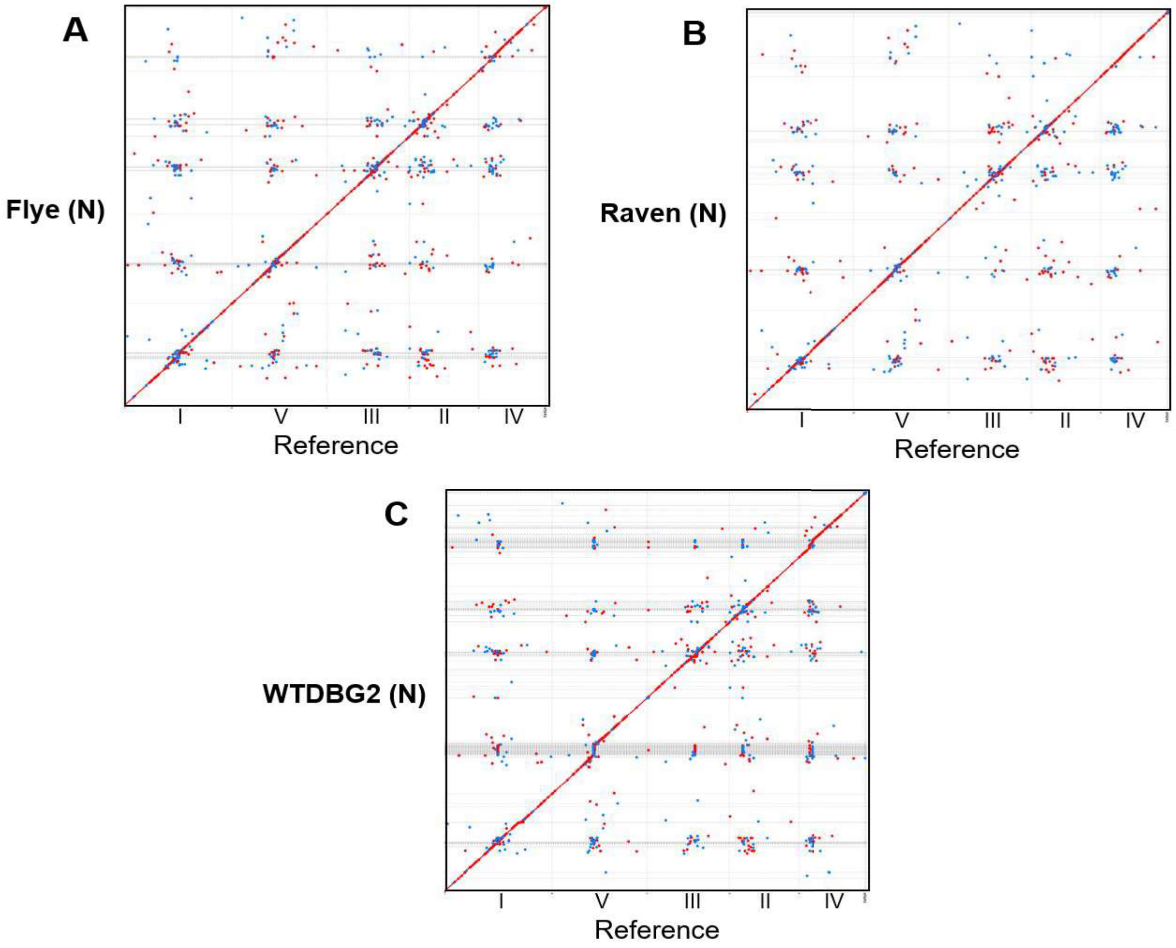


**Supplementary Figure S2 |** Dot plots for all final Oxford Nanopore candidate assemblies generated for *A. thaliana* with the *Pyro* pipeline, as generated by Mashmap in comparison to the TAIR10 reference assembly. (I) = Illumina, (N) = Nanopore, (P) = PacBio, (H) = Hybrid. All assemblies show high completeness when compared with the reference, and considerably higher contiguity versus the Illumina assemblies in **Supplementary Figure S1**. There is some noticeable variation in the length of the centromeric regions, such as of chromosome 5 and chromosome 4, particularly by WTDBG2, with notably higher contiguity and lower fragmentation in such regions in the Raven assembly.

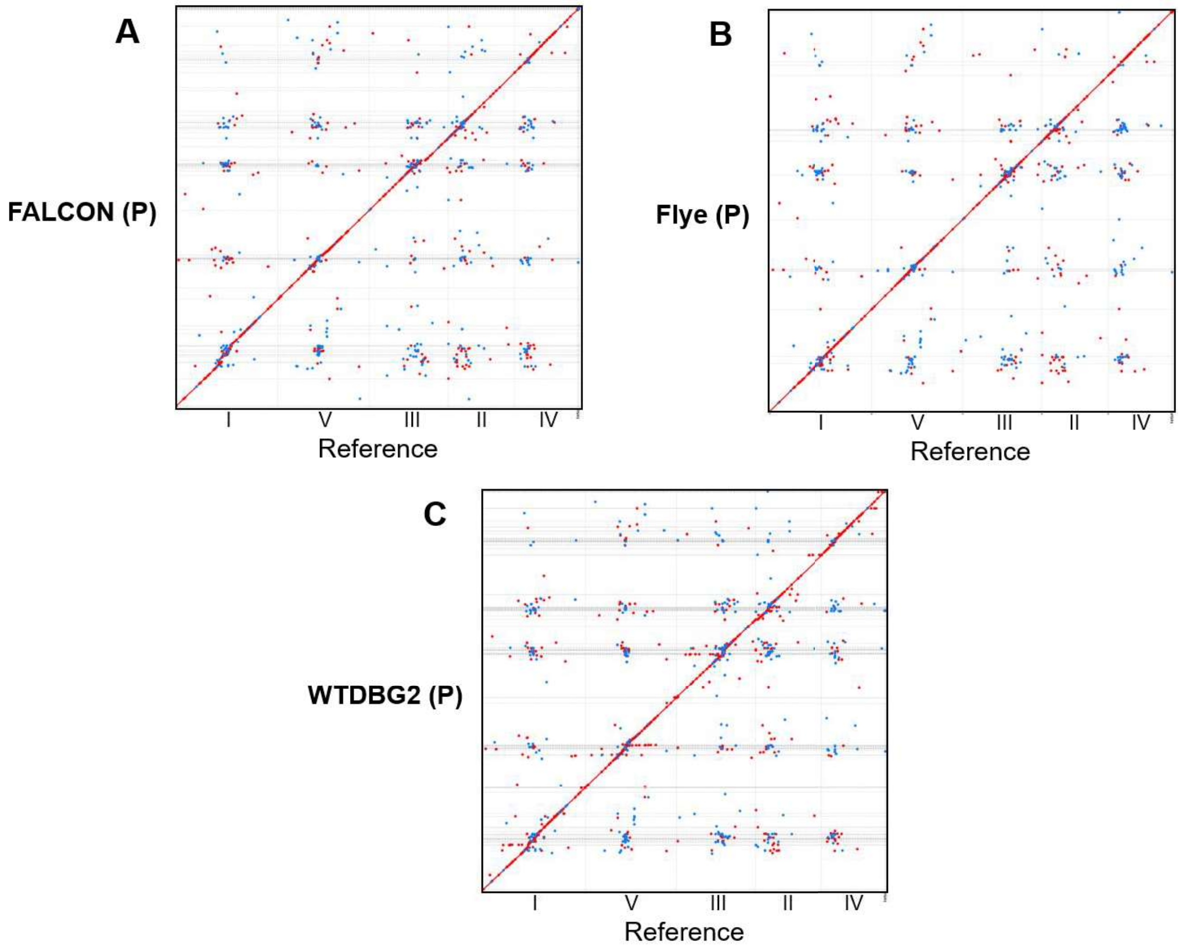


**Supplementary Figure S3 |** Dot plots for all final PacBio CLR candidate assemblies generated for *A. thaliana* with the *Pyro* pipeline, as generated by Mashmap in comparison to the TAIR10 reference assembly. (I) = Illumina, (N) = Nanopore, (P) = PacBio, (H) = Hybrid. All assemblies show high completeness when compared with the reference, and comparable contiguity to the Oxford Nanopore assemblies in **Supplementary Figure S2**. Again, there is some noticeable disagreement in the lengths of the centromeric regions between assemblers, in particular for chromosome 1.

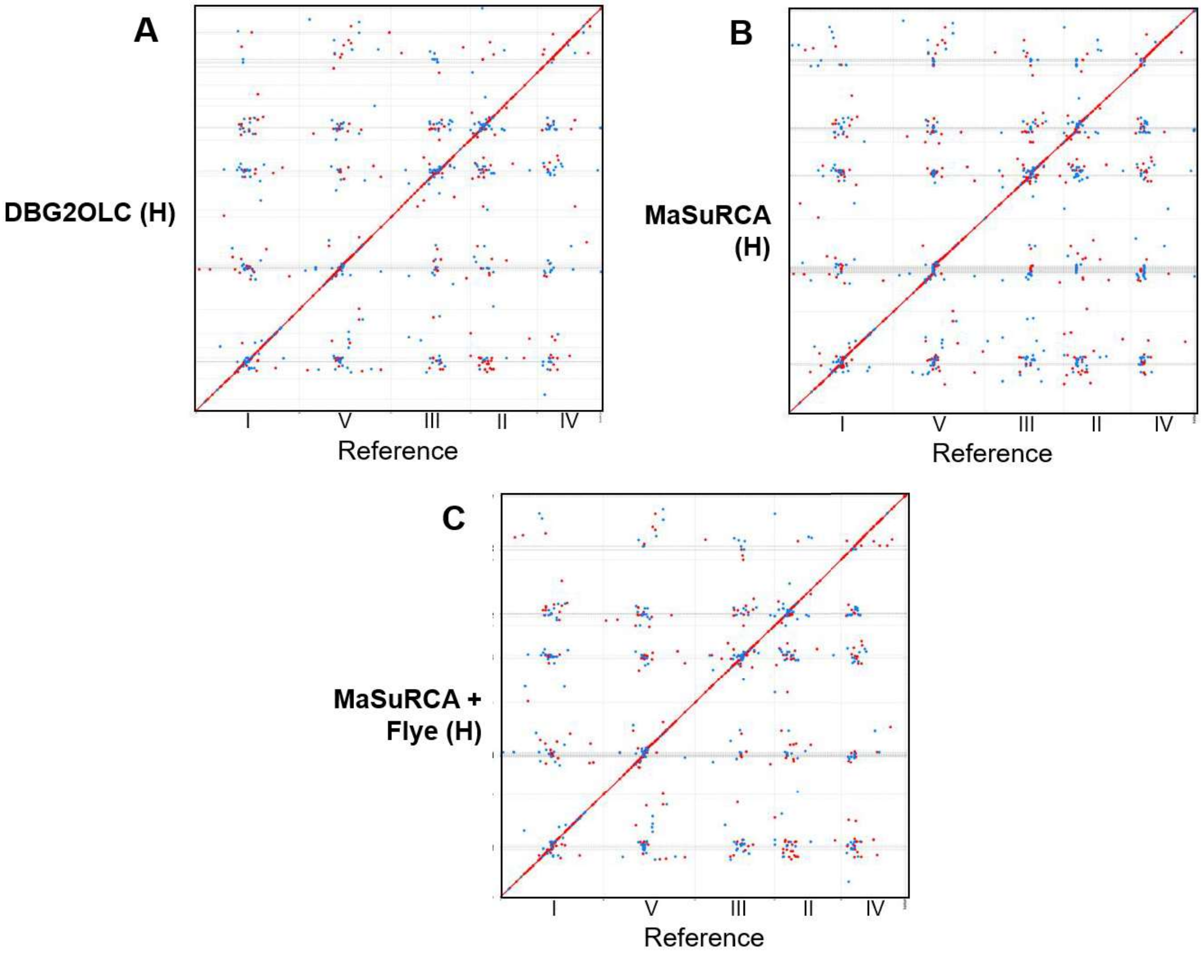


**Supplementary Figure S4 |** Dot plots for all final hybrid candidate assemblies generated for *A. thaliana* with the *Pyro* pipeline, as generated by Mashmap in comparison to the TAIR10 reference assembly. (I) = Illumina, (N) = Nanopore, (P) = PacBio, (H) = Hybrid. All assemblies show high completeness when compared with the reference, and comparable contiguity to the Oxford Nanopore and PacBio CLR assemblies in **Supplementary Figures S2–S3**. As above, there is some disagreement in the lengths of centromeric regions, again around chromosome 5, particularly for the MaSuRCA assembly.

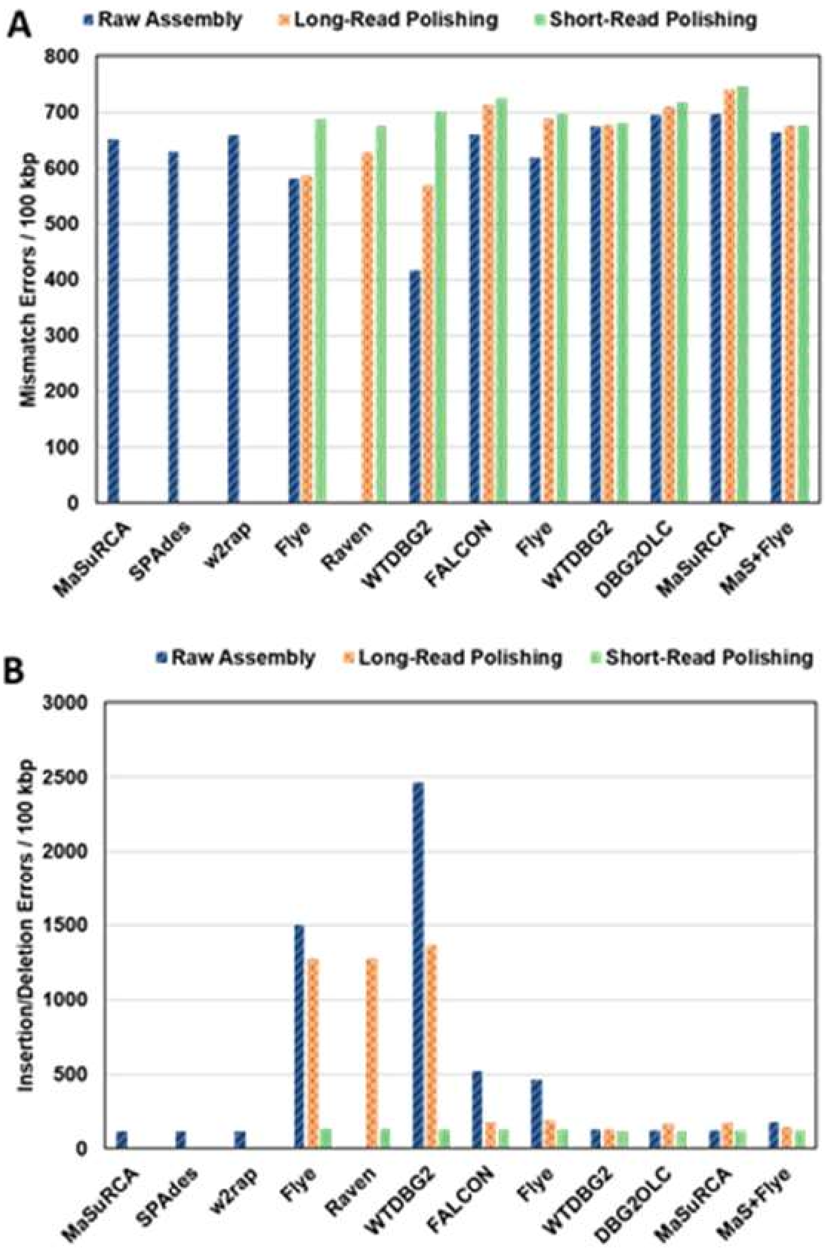


**Supplementary Figure S5 |** Comparisons of mismatch errors and insertion/deletion errors across sequencing types and polishing methods, as calculated by QUAST v 5.0.2 in comparison with the TAIR10 reference assembly, determined after each major stage of the pipeline, namely after assembly, after long-read polishing, and after short-read polishing. Illumina-only assembly methods were not polished, and so have no values for long-read or short-read polishing. The initial raw Raven assembly was undetectable by QUAST, and therefore has no values, but indicates large but undeterminable error rates. **(A)** Mismatch errors across assemblers and polishing methods. Initial lower values for Oxford Nanopore assemblies may be due to regions containing further mismatches being undetectable before some initial polishing. **(B)** Insertion/deletion errors across assemblers and polishing methods. Errors are in particular lowered considerably with the addition of short-read polishing for Oxford Nanopore assemblies, offsetting the comparably low increase in mismatch errors, and highlighting the value of utilizing short and long reads for sequencing projects.

#### 1.2 Supplementary Tables

**Supplementary Table S1.** Full details of the third-party packages used in the *Prep* module of the *Pyro* pipeline for data preparation. Current versions employed in the pipeline are given, along with availability and the default *Pyro* commands for packages where package defaults are not used – these can be changed by user input *via* the config file.

| Package | Version | Citation | Description |
|---|---|---|---|
| Trim_Galore | 0.6.4 | - | Adapter trimming and quality filtering of Illumina PE data. |
| | https://www.bioinformatics.babraham.ac.uk/projects/trim_galore/ | | |
| | trim_galore --cores 4 --quality 20 --length 90 --trim-n --max_n 0 --paired {reads} | | |
| FiltLong | 0.2.0 | - | Quality filtering and processing of long-read data. |
| | https://github.com/rrwick/Filtlong | | |
| | filtlong --min_length 1000 --keep_percent 90 {reads} | | |
| BBMap | 38.86 | Bushnell, 2014 | Random subsampling of input data based on desired coverage level and estimated genome size. |
| | https://sourceforge.net/projects/bbmap/ | | |
| QuorUM | 1.1.1 | Marçais et al., 2015 | Error correction of Illumina PE . reads |
| | https://github.com/alekseyzimin/masurca | | |
| Canu | 2.0 | Koren et al., 2017 | Error correction of long-read data. |
| | https://github.com/marbl/canu | | |
| Ratatosk | 0.3.0 | Holley et al., 2020 | Error correction of Oxford Nanopore reads using Illumina PE data. |
| | https://github.com/DecodeGenetics/Ratatosk | | |

**Supplementary Table S2.** Full details of the third-party packages used in the *Build* module of the *Pyro* pipeline for draft assembly construction. Current versions employed in the pipeline are given, along with availability and the default commands and parameters set for all packages, unless changed by user input *via* the config file.

| Package | Version | Citation | Description |
|---|---|---|---|
| ABySS | 2.1.0 | Jackman et al., 2017 | Short-read de Bruijn graph-based assembly and scaffolding package using Bloom filters. Authors emphasize scalability and parallelizability for large genomes |
| | https://github.com/bcgsc/abyss | | |
| | abyss-pe name={prefix} k=96 in='{reads}' j={threads} B=2G H=4 kc=2 v=-v | | |
| MaSuRCA | 3.4.1 | Zimin et al., 2013 | Short-read and hybrid assembler using a mix of de Bruijn graph and overlap-layout-consensus approaches. High accuracy, but can require more time commitment due to error correction steps. |
| | https://github.com/alekseyzimin/masurca | | |
| | masurca {config};<br>bash assemble.sh;<br>[uses default config file – please see https://github.com/genomeassembler/pyro for default] | | |
| Meraculous | 2.2.5.1 | Chapman et al., 2011 | Short-read assembly package using de Bruijn graphs. Usually requires >30X coverage to run as intended. |
| | https://sourceforge.net/projects/meraculous20/ | | |
| | run_meraculous.sh -c {config} -dir {workdir} -cleanup_level=1<br>[uses config file – please see https://github.com/genomeassembler/pyro for default] | | |
| Platanus | 1.2.4 | Kajitani et al., 2014 | Short-read assembly and scaffolding pipeline using de Bruijn graphs. Also usually requires >30X coverage to run as intended |
| | http://platanus.bio.titech.ac.jp/platanus-assembler/platanus-1-2-4 | | |
| | platanus assemble -o {prefix} -f {reads} -k 32 -t {threads} -m {memory};<br>platanus scaffold -o {prefix} -c {contigs} -b {bubbles} -IP1 {reads} -t {threads}; | | |

| | | | |
|---|---|---|---|
| | platanus gap_close -o {prefix} -c {scaffolds} -f {reads} -t {threads}; | | |
| Package | Version | Citation | Description |
| Ray | 2.3.1 | Boisvert et al., 2010 | Short-read assembly package using de Bruijn graphs. Fairly robust and reliable, but can require higher time commitments than some other assemblers. |
| | https://github.com/sebhtml/ray | | |
| | mpiexec -n {threads} Ray -p {reads} -o {outdir} | | |
| SOAP-denovo2 | r240 | Luo et al., 2012 | Short-read assembly and scaffolding package using sparse de Bruijn graphs. Performance tends to rely more heavily on defining *k*-mer length than other assemblers. |
| | https://github.com/aquaskyline/SOAPdenovo2 | | |
| | SOAPdenovo-63mer all -s {config} -o {outdir} -K 51 -p {threads} -N {genome-size}<br>[uses config file – please see https://github.com/genomeassembler/pyro for default] | | |
| SPAdes | 3.12.0 | Bankevich et al., 2012 | Short-read and hybrid assembly package using de Bruijn graphs. Can require more time than other assembly packages due to error correction of reads, but also produces consistently high quality assemblies. |
| | https://github.com/ablab/spades | | |
| | spades.py -1 {reads} -2 {reads} -t {threads} -m {memory} -o {outdir} | | |
| Sparse-Assembler | 20160920 | Ye et al., 2012 | Short-read assembly package using sparse de Bruijn graphs. Memory efficient, but not parallelized, only single core. Useful to pair with DBG2OLC for scaffolding of contigs, or hybrid assembly. |
| | https://github.com/yechengxi/SparseAssembler | | |
| | SparseAssembler g 10 k 51 LD 0 GS {genome-size} f {reads} f {reads};<br>DBG2OLC LD1 0 Contigs {contigs} k 31 KmerCovTh 0 MinOverlap 50 PathCovTh 1 f {reads} f {reads}; | | |

| Package | Version | Citation | Description |
| --- | --- | --- | --- |
| w2rap | 20180828 | Clavijo et al., 2017 | Short-read assembly and scaffolding pipeline using de Bruijn graphs, improving upon on the DISCOVAR de novo pipeline. Fast but can be memory-intensive for large genomes |
| | https://github.com/bioinfologics/w2rap-contigger | | |
| | w2rap-contigger -t {threads} -m {memory} -o {outdir} -r {reads} -p {prefix} | | |
| Canu | 1.9 | Koren et al., 2017 | Long-read error correction and assembly pipeline using an overlap-layout-consensus approach. Error correction takes significant resources, particularly for Oxford Nanopore inputs, but can exploit computing grids for speed-up. Produces high accuracy, high contiguity assemblies but requires considerable tuning and configuration for best results. |
| | https://github.com/marbl/canu | | |
| | canu -p {prefix} -d {outdir} genomeSize={genome-size} –{pacbio or nanopore}-raw {reads} -useGrid=false; | | |
| FALCON | 0.3.0 | Chin et al., 2016 | Long-read assembly package specifically for PacBio reads. Can require increased time and resources when configuration is not optimized. |
| | https://github.com/PacificBiosciences/FALCON | | |
| | fc_run falcon.cfg<br>[uses config file – please see https://github.com/genomeassembler/pyro for default] | | |
| Flye | 2.8 | Kolmogorov et al., 2019 | Long-read assembly package based on repeat graphs. Middle-ground resource usage, but consistently high contiguity and accuracy. |
| | https://github.com/fenderglass/Flye | | |
| | flye --{nanopore or pacbio}-raw {reads} --out-dir {outdir} --threads {threads} | | |

| Package | Version | Citation | Description |
|---|---|---|---|
| MECAT2 | 20190314 | Xiao et al., 2017 | Long-read assembly package specifically for PacBio reads. Middle-ground resource usage, typically requires higher coverage (40X +) for contiguous and complete assembly. |
| | https://github.com/xiaochuanle/MECAT2 | | |
| | mecat.pl correct {config};<br>mecat.pl trim {config};<br>mecat.pl assemble {config};<br>[uses config file – please see https://github.com/genomeassembler/pyro for default] | | |
| Miniasm | 0.3-r179 | Li, 2016 | Long-read assembly package using overlap-layout, without consensus step. Fast, but raw assembly requires polishing to be of use. |
| | https://github.com/lh3/miniasm | | |
| | minimap2 -x ava-{ont or pb} -t {threads} {reads} {reads};<br>miniasm -f {reads} {align}; | | |
| NECAT | 0.0.1_ update 20200803 | | Long-read assembly and scaffolding package specifically for Oxford Nanopore reads. Middle-ground resource usage, but can run on low coverage (20X) and still perform well. |
| | https://github.com/xiaochuanle/NECAT | | |
| | necat.pl bridge {config};<br>[uses config file – please see https://github.com/genomeassembler/pyro for default] | | |
| Raven | 0.0.5 | Vaser and Sikic, 2020 | Long-read assembly package for uncorrected reads. Fast, and consistently produces highly contiguous assemblies. Raw assembly requires polishing to be useful. |
| | https://github.com/lbcb-sci/raven | | |
| | raven -p 0 -t {threads} {reads} | | |

| Package | Version | Citation | Description |
| --- | --- | --- | --- |
| Shasta | 0.3.0 | Shafin et al., 2020 | Long-read assembly package specifically for Oxford Nanopore reads. Very fast, low resource usage, but tends to require high coverage (> 40X) to guarantee success across the genome. |
| | https://github.com/chanzuckerberg/shasta | | |
| | shasta --input {reads} --threads {threads} | | |
| WTDBG2 | 2.5 | Ruan and Li, 2020 | Long-read assembly package using de Bruijn graphs. Fast and light on memory usage while maintaining contiguity, usually requires polishing to be useful for accuracy. |
| | https://github.com/ruanjue/wtdbg2 | | |
| | wtdbg2 -x {ont or rs} -g {genome-size} -t {threads} -i {reads} -fo {prefix};<br>wtpoa-cns -t {threads} -i {contigs} -fo {outfile}; | | |
| DBG2OLC | 20160920 | Ye et al., 2016 | Hybrid assembly pipeline which takes PacBio reads and a short-read assembly as inputs. Significant parts of the pipeline are single-core, so tends to require more time to run. Runs better for low coverage of long reads (15-20X), high contiguity and accuracy. By default, runs SparseAssembler for Illumina contig input, and runs Sparc for fixing output assembly. |
| | https://github.com/yechengxi/DBG2OLC | | |
| | SparseAssembler g 10 k 51 LD 0 GS {genome-size} f {reads} f {reads};<br>DBG2OLC Contigs {contigs} k 17 KmerCovTh 2 MinOverlap 50 AdaptiveTh 0.01 RemoveChimera 1 f {reads};<br>cat {contigs} {reads} > {all};<br>split_reads_by_backbone.py -b {backbone} -o {workdir} -r {all} -c {consensus-info};<br>split_and_run_sparc.sh {backbone} {consensus-info} {all} {workdir} 2 > {log}; | | |

| Package | Version | Citation | Description |
| --- | --- | --- | --- |
| HASLR | 0.8a1 | Haghshenas et al., 2020 | Hybrid short- and long-read assembler. Lightning fast with minimal resource requirements. High contiguity, and provides higher completeness than most raw long-read assemblies without polishing, but can fall behind when polishing is taken into account. |
| | https://github.com/vpc-ccg/haslr | | |
| | python haslr.py -t {threads} -o {outdir} -g {genome-size} -l {longreads} -x {nanopore or pacbio} -s {shortreads} | | |

**Supplementary Table S3.** Full details of the third-party packages used in the *Fix* module of the *Pyro* pipeline for assembly polishing. Current versions employed in the pipeline are given, along with the descriptions of each package. All packages are run with baseline default parameters.

| Package | Version | Citation | Description |
|---|---|---|---|
| HyPo | 1.0.3 | Kundu et al., 2019 | Short-read or hybrid short- and long-read polishing package. Fast and consistent in improving assembly completeness. |
| | https://github.com/kensung-lab/hypo | | |
| NextPolish | 1.3.0 | Hu et al., 2020 | Short- and long-read polishing package. Can be resource-intensive, but consistently produces substantial improvement in assembly quality whether run with short- or long-reads. |
| | https://github.com/Nextomics/NextPolish | | |
| ntEdit | 1.3.2 | Warren et al., 2019 | Short-read polishing package. Very fast, but considerably more conservative in fixing errors as other packages. Prior setup is based on reads, so all improvement is within one round. |
| | https://github.com/bcgsc/ntEdit | | |
| Pilon | 1.23 | Walker et al., 2014 | Short-read polishing package. Careful and reliable, but slower than other short-read polishing options. Can take two to three rounds to reach highest possible quality. |
| | https://github.com/broadinstitute/pilon/ | | |
| POLCA | 3.4.1 | Zimin and Salzberg, 2020 | Short-read polishing package. Fast and conservative in changes made, but consistently improves assembly quality. |
| | https://github.com/alekseyzimin/masurca | | |
| Racon | 1.4.13 | Vaser et al., 2017 | Short- and long-read polishing package. Tends to be less conservative than other polishing packages, but it is flexible, and can produce the highest quality results out of the options available. |
| | https://github.com/lbcb-sci/racon | | |

| Package | Version | Citation | Description |
|---|---|---|---|
| Arrow | 0.3.0 | Chin et al., 2016 | Long-read polishing package specifically using PacBio reads. Requires fresh-off-the-sequencer PacBio data with correct feature headers/formats to work. Can produce high quality results when conditions are satisfied, will not run if they are not. |
| | https://github.com/PacificBiosciences/GenomicConsensus | | |
| Medaka | 0.12.1 | - | Long-read polishing package specifically for Oxford Nanopore reads. Tends to take longer than other options to run, but produces comparably high quality. |
| | https://github.com/nanoporetech/medaka | | |

**Supplementary Table S4.** Full details of the third-party packages used in the *Check* module of the *Pyro* pipeline for quality metric evaluation and data presentation. Current versions employed in the pipeline are given – all packages are run with baseline default parameters, unless otherwise noted below the entry.

| Package | Version | Citation | Description |
|---|---|---|---|
| assembly-stats | 1.0.1 | - | Very light package which calculates basic assembly statistics such as N50, average length, total assembly size, and number of fragments. |
| | https://github.com/sanger-pathogens/assembly-stats | | |
| BUSCO | 3.1.0 | Seppey et al., 2019 | Package for calculating gene completeness of an assembly using BUSCOs. Requires an OrthoDB dataset to be specified to run. |
| | https://gitlab.com/ezlab/busco/ | | |
| BUSCOMP | 0.9.2 | Edwards, 2019 | Package to calculate more thorough comparative analysis of BUSCO results across assemblies. Requires BUSCO results to run. |
| | https://github.com/slimsuite/buscomp | | |
| FastQC | 0.11.8 | - | Package to assess the quality of sequencing reads, both pre- and post-processing in terms of metrics such as GC percentage, length distributions, and the presence of adapter sequences. |
| | https://www.bioinformatics.babraham.ac.uk/projects/fastqc/ | | |
| Jellyfish | 2.3.0 | Garçais and Kingsford, 2011 | Package to calculate *k*-mer distributions based on short-read sequencing. Creates necessary input for GenomeScope for estimating genome size. |
| | https://github.com/gmarcais/Jellyfish | | |
| GenomeScope | 1.0.0 | Vurture et al., 2017 | Package to estimate genome size and heterozygosity from *k*-mer distributions calculated by Jellyfish. |
| | https://github.com/schatzlab/genomescope | | |

| Package | Version | Citation | Description |
| --- | --- | --- | --- |
| Mashmap | 2.0 | Jain et al., 2018a | Package to align reference sequences to draft assemblies to create dot plots. |
| | https://github.com/marbl/MashMap | | |
| QUAST | 5.0.2 | Mikheenko et al., 2018 | Package to calculate reference-based statistics such as mismatches, misassemblies, insertion/deletion errors, and genome fraction. |
| | http://quast.sourceforge.net/ | | |
| | python quast.py -o {outdir} -r {reference} -t {threads} -m 200 --extensive-mis-size 7000 --min-contig 3000 --min-alignment 500 --space-efficient --eukaryote --fast -l {labels} -1 {shortreads} -2 {shortreads} --{pacbio or nanopore} {longreads} {assemblies} | | |
| Red | 2.0 | Girgis, 2015 | Package to calculate overall repeat percentage for assemblies. Faster than other options, but provides less detailed breakdown. |
| | https://github.com/BioinformaticsToolsmith/Red | | |

**Supplementary Table S5.** Full results for all major assemblies generated, including raw assemblies and final steps of each method of polishing, highlighting the contiguity, gene completeness and accuracy of all assemblies generated, as well as the value of fully automated, fully integrated polishing of assemblies with both long and short reads for heightened assembly quality. Values in bold are the 'best' among all *Pyro* assemblies for metrics where it is useful to rank based on value – that is, highest for N50, BUSCO Complete, BUSCOMP Complete, and BUSCOMP Identity, and lowest for Fragments, Misassemblies, Mismatches / 100 kbp and Insertions or Deletions / 100 kbp.

| **Assembler** | **Polishing Strategies** | **Assembly Statistics and Contiguity** | | | | **Gene Completeness and Identity** | | | **Reference-Based Metrics** | | |
|---|---|---|---|---|---|---|---|---|---|---|---|
| | | **Assembly Size (Mbp)** | **Fragments** | **N50 (Mbp)** | **Total Repeat Length (Mbp)** | **BUSCO Complete (%)** | **BUSCOMP Complete (%)** | **BUSCOMP Identity (%)** | **Mis-assemblies** | **Mismatches / 100 kbp** | **Insertions or Deletions / 100 kbp** |
| **Illumina Paired-End** | | | | | | | | | | | |
| MaSuRCA | - | 122.4 | 8,689 | 0.400 | 43.8 | **98.4** | 99.8 | 68.3 | 1788 | 657.7 | 118.5 |
| SPAdes | - | 126.7 | 49,791 | 0.228 | 43.2 | 98.3 | 99.6 | 68.3 | 1173 | 628.3 | **117.0** |
| w2rap | - | 136.4 | 58,337 | 0.255 | 40.4 | **98.4** | 99.7 | 67.6 | 1474 | 650.9 | 118.2 |
| **Oxford Nanopore** | | | | | | | | | | | |
| Flye | - | 120.4 | 118 | 14.073 | 47.6 | 36.4 | **99.9** | 0.1 | 810 | 581.1 | 1501.2 |
| | NextPolishx4 (ONT) | 120.7 | 118 | 14.111 | 48.2 | 45.3 | **99.9** | 0.8 | 887 | 585.7 | 1274.5 |
| | HyPo x3 (ILL) | 122.8 | 118 | 14.358 | 44.3 | 98.3 | **99.9** | 68.4 | 1922 | 688.1 | 132.6 |
| Raven | - | 110.8 | **32** | 10.047 | 60.4 | 0.4 | 88.2 | 0.0 | - | - | - |
| | NextPolish x4 (ONT) | 117.7 | **32** | 10.674 | 49.9 | 46.4 | **99.9** | 0.8 | 764 | 627.5 | 1271.0 |
| | HyPo x3 (ILL) | 119.7 | **32** | 10.850 | 50.3 | 98.3 | **99.9** | 68.5 | 1929 | 675.4 | 131 |

| Assembler | Polishing Strategies | Assembly Statistics and Contiguity | | | | Gene Completeness and Identity | | | Reference-Based Metrics | | |
|---|---|---|---|---|---|---|---|---|---|---|---|
| | | **Assembly Size (Mbp)** | **Fragments** | **N50 (Mbp)** | **Total Repeat Length (Mbp)** | **BUSCO Complete (%)** | **BUSCOMP Complete (%)** | **BUSCOMP Identity (%)** | **Mis-assemblies** | **Mismatches / 100 kbp** | **Insertions or Deletions / 100 kbp** |
| **Oxford Nanopore** | | | | | | | | | | | |
| WTDBG2 | - | 118.1 | 372 | 3.739 | 52.7 | 14.5 | 98.5 | 0.0 | **202** | **416.6** | 2460.5 |
| | NextPolish x4 (ONT) | 121.2 | 372 | 3.806 | 46.1 | 41.5 | 98.8 | 0.5 | 712 | 568.8 | 1367.5 |
| | HyPo x3 (ILL) | 123.4 | 372 | 3.875 | 46.8 | 97.4 | 98.8 | 67.5 | 1865 | 701.3 | 131.0 |
| **PacBio CLR** | | | | | | | | | | | |
| FALCON | - | 121.0 | 122 | 5.042 | 43.2 | 83.3 | **99.9** | 18.1 | 1697 | 618.2 | 465.7 |
| | NextPolish x4 (PB) | 121.3 | 122 | 5.054 | 43.6 | 95.6 | **99.9** | 54.5 | 1977 | 688.5 | 191.0 |
| | POLCA x3 (ILL) | 121.4 | 122 | 5.057 | 43.7 | 98.2 | **99.9** | 67.4 | 2030 | 696.9 | 126.8 |
| Flye | - | 120.3 | 76 | 14.331 | 43.7 | 98.2 | **99.9** | 67.4 | 1906 | 674.5 | 130.7 |
| | NextPolish x4 (PB) | 120.3 | 76 | 14.330 | 43.9 | 97.9 | **99.9** | 67.5 | 1915 | 677.9 | 130.1 |
| | POLCA x3 (ILL) | 120.3 | 76 | 14.332 | 43.9 | 98.3 | **99.9** | 68.4 | 1918 | 681.0 | 119.8 |
| WTDBG2 | - | 123.7 | 312 | 9.979 | 48.9 | 81.1 | 99.6 | 20.0 | 1456 | 659.0 | 521.7 |
| | NextPolish x4 (PB) | 124.0 | 312 | 9.996 | 46.1 | 96.5 | **99.9** | 58.7 | 1888 | 713.1 | 178.8 |
| | POLCA x3 (ILL) | 124.0 | 312 | 10.001 | 46.0 | 98.3 | **99.9** | 68.3 | 1949 | 723.7 | 125.7 |

| Assembler | Polishing Strategies | Assembly Statistics and Contiguity | | | | Gene Completeness and Identity | | | Reference-Based Metrics | | |
|---|---|---|---|---|---|---|---|---|---|---|---|
| | | Assembly Size (Mbp) | Fragments | N50 (Mbp) | Total Repeat Length (Mbp) | BUSCO Complete (%) | BUSCOMP Complete (%) | BUSCOMP Identity (%) | Mis-assemblies | Mismatches / 100 kbp | Insertions or Deletions / 100 kbp |
| **Hybrid assemblies** | | | | | | | | | | | |
| DBG2OLC | - | 119.3 | 65 | 6.923 | 45.7 | 95.9 | 99.6 | **68.7** | 1848 | 663.4 | 177.8 |
| | NextPolish x4 (PB) | 119.2 | 65 | 6.916 | 49.9 | 97.5 | 99.6 | 67.9 | 1942 | 675.1 | 143.3 |
| | HyPo x3 (ILL) | 119.2 | 65 | 6.914 | 49.8 | 98.2 | 99.6 | 68.1 | 1981 | 675.6 | 122.3 |
| MaSuRCA | - | 126.8 | 105 | 14.486 | 44.4 | 98.1 | **99.9** | 68.3 | 2159 | 696.9 | 121.4 |
| | NextPolish x4 (ONT+PB) | 126.8 | 105 | 14.481 | 41.9 | 97.5 | **99.9** | 61.9 | 2145 | 740.3 | 173.4 |
| | HyPo x3 (ILL) | 126.9 | 105 | 14.488 | 41.7 | 98.3 | **99.9** | 68.3 | 2186 | 745.9 | 122.8 |
| MaSuRCA + Flye | - | 121.9 | 124 | 14.865 | 43.4 | 98.2 | **99.9** | 68.4 | 2005 | 694.5 | 120.2 |
| | NextPolish x4 (ONT+PB) | 121.9 | 124 | 14.860 | 44.3 | 97.5 | **99.9** | 61.7 | 1983 | 709.7 | 167.7 |
| | HyPo x3 (ILL) | 122.0 | 124 | **14.867** | 44.0 | 98.3 | **99.9** | 68.4 | 2030 | 717.2 | 119.9 |
| **Reference** | | | | | | | | | | | |
| - | - | 119.7 | 7 | 23.460 | 43.9 | 98.2 | 99.8 | 51.9 | - | - | - |

**Supplementary Table S6 |** Computational performance of each assembly strategy for the *A. thaliana* dataset. Each assembler in the *Pyro* pipeline was run on a single 128 Gb RAM node with 20 CPUs. While the *Pyro* pipeline recommends a combination of the most efficient and most effective assemblers given a certain dataset, there is often a trade-off between assembly quality and computational performance. Therefore, there is a more significant time burden for some sequencing strategies, such as hybrid assemblers.

| Sequencing Technology | Assembly Package | Assembly | | Long-Read Polishing | | Short-Read Polishing | | Total Time to Final Assembly |
|---|---|---|---|---|---|---|---|---|
| | | Wall Clock Time (hh:mm:ss) | Peak Memory Usage (Gb) | Wall Clock Time (hh:mm:ss) | Peak Memory Usage (Gb) | Wall Clock Time (hh:mm:ss) | Peak Memory Usage (Gb) | |
| **Illumina** | MaSuRCA | 3:03:34 | 23.7 | - | - | - | - | 3:03:34 |
| | SPAdes | 5:52:15 | 49.6 | - | - | - | - | 5:52:15 |
| | w2rap | 0:35:43 | 94.9 | - | - | - | - | 0:35:43 |
| **Oxford Nanopore** | Flye | 1:23:07 | 30.1 | 0:50:37 | 32.4 | 1:32:53 | 17.9 | 3:46:37 |
| | Raven | 0:12:52 | 20.0 | 0:55:20 | 46.0 | 1:28:20 | 21.5 | 2:36:32 |
| | WTDBG2 | 0:23:10 | 14.9 | 0:49:29 | 33.5 | 1:30:17 | 18.1 | 2:42:56 |
| **PacBio CLR** | FALCON | 7:43:59 | 112.5 | 1:41:10 | 74.4 | 2:18:09 | 19.9 | 11:43:18 |
| | Flye | 2:08:32 | 37.0 | 1:37:09 | 55.9 | 2:59:07 | 19.3 | 6:44:48 |
| | WTDBG2 | 0:18:34 | 13.6 | 1:46:04 | 46.4 | 2:21:23 | 19.3 | 4:26:01 |
| **Hybrid** | DBG2OLC | 4:21:32 | 7.6 | 1:33:30 | 73.4 | 1:26:42 | 18.0 | 7:21:44 |
| | MaSuRCA | 13:07:51 | 34.9 | 2:21:41 | 54.2 | 1:25:02 | 18.1 | 16:54:34 |
| | MaSuRCA+Flye | 8:16:03 | 27.5 | 2:17:14 | 54.6 | 1:31:27 | 18.1 | 12:04:44 |
